# Identification of bazedoxifene for the treatment of LGMD R2 by high throughput screening

**DOI:** 10.1101/2024.02.28.582550

**Authors:** Celine Bruge, Nathalie Bourg, Emilie Pellier, Johana Tournois, Jerome Polentes, Manon Benabides, Noella Grossi, Anne Bigot, Anthony Brureau, Isabelle Richard, Xavier Nissan

**Affiliations:** Université Paris-Saclay, Université d’Evry, Inserm, IStem, UMR861, Corbeil-Essonnes, 91100 France; IStem, CECS, Corbeil-Essonnes, 91100, France; INTEGRARE, Genethon, Inserm, Univ Evry, Université Paris-Saclay, Evry, 91002, France; Sorbonne Université, Inserm, Institut de Myologie, Centre de Recherche en Myologie, Paris, 75013, France

**Keywords:** limb-girdle muscular dystrophies, misfolded protein, high-throughput screening, saracatinib, bazedoxifene, autophagy

## Abstract

LGMD R2 is a rare genetic disorder characterized by progressive proximal muscle weakness and wasting caused by a recessive loss of function of dysferlin, a transmembrane protein controlling plasma membrane repair in skeletal muscles. We report here the development of an *in vitro* high-throughput assay using immortalized myoblasts and monitored reallocation of an aggregated mutant form of dysferlin (*DYSF*^L1341P^). Using this assay, we screened a library of 2239 drugs and identified two autophagy inducers, namely saracatinib and bazedoxifene, as potential drugs to repurpose for LGMD R2 patients carrying the *DYSF*^L1341P^ mutation. Functional characterization of these drugs revealed that saracatinib and bazedoxifene had a protective effect on the plasma membrane in osmotic shock assay. While saracatinib restores functionality in membrane resealing through a specific rescue of L1341P dysferlin from degradation, bazedoxifene demonstrates an additional protective effect on dysferlin KO mice muscle fibers. Finally, further investigations into the molecular mechanism of action of bazedoxifene revealed an induction of autophagy flux, which may underlie the molecule’s effect on the survival of LGMD R2 myofibers.

## INTRODUCTION

Limb-girdle muscular dystrophies (LGMD) refer to a genetically heterogeneous group of autosomal inherited muscular dystrophies, characterized by progressive muscle weakness and wasting of the shoulder or pelvic girdles (Walton & Nattrass, 1954). Among LGMD, dysferlinopathies are a subgroup caused by mutations in the dysferlin gene (*DYSF*) (Bashir *et al*, 1998; Liu *et al*, 1998) and are mainly defined by the two most frequently encountered clinical phenotypes (Nguyen *et al*, 2007): distal Miyoshi myopathy type 1 (MM) and proximal LGMD R2 (previously named 2B). Dysferlinopathies are characterized by phenotypic heterogeneity with a high degree of variability in disease onset, disease severity and progression (LoMauro *et al*, 2021). The protein encoded by the *DYSF* gene, dysferlin, is a type-II transmembrane protein mainly expressed in the sarcolemma of striated skeletal muscles (Anderson *et al*, 1999). Dysferlin belongs to the ferlin family and contains a large intracellular domain composed of seven calcium-binding C2-domains, a carboxy-terminal transmembrane region and a very short extracellular domain. While the clinical features of dysferlinopathies are well described, the molecular mechanisms related to these diseases are still not fully characterized. Over recent years, studies on dysferlin have mostly focused on its role in membrane repair of injured sarcolemma in skeletal muscle fibers; a critical calcium-dependent machinery for maintaining muscle integrity and function (Blazek *et al*, 2015). Ultrastructural analysis of dysferlinopathy muscle fibers showed disruption of the plasma membrane and its replacement by layers of small vesicles (Selcen *et al*, 2001; Cenacchi *et al*, 2005). Loss of membrane integrity and accumulation of subsarcolemmal vesicles were also observed in muscle fibers from dysferlin-deficient mice (Bansal *et al*, 2003). Other suggested functions of dysferlin, that could contribute toward the pathophysiology of dysferlinopathies, include vesicular trafficking (Bansal *et al*, 2003; Lek *et al*, 2012), monocyte adhesion (de Morrée *et al*, 2013), inflammation (Han, 2011; McNally *et al*, 2000) and calcium homeostasis (Kerr *et al*, 2014; Cea *et al*, 2016). Recent evidence indicates that dysferlin may also be involved in the regulation of the T-tubule system (Demonbreun *et al*, 2014; Kerr *et al*, 2013; Klinge *et al*, 2007, 2010), lipid metabolism and fat deposition (Grounds *et al*, 2014; Haynes *et al*, 2019). While several therapeutic strategies using gene therapy or exon skipping have been investigated (Barthélémy *et al*, 2018; Lee *et al*, 2018; Lostal *et al*, 2010; Potter *et al*, 2018; Pryadkina *et al*, 2015; Verwey *et al*, 2020), there is currently no treatment for LGMD R2. Recent studies highlight the potential of targeting pathological phenotypes, as revealed through the use of galectin-1 (Gal-1, a soluble carbohydrate-binding protein), to increase the membrane repair capacity and myogenic potential of dysferlin-deficient muscle cells and muscle fibers (Vallecillo-Zúniga *et al*, 2020). Identification of additional pharmacologic compounds that would decrease the course of the disease remains consequently a major strategy for LGMD R2 therapy.

Analyses of large cohorts of patients have revealed that several hundred mutations in the *DYSF* gene can lead to dysferlinopathies (Leiden Open Variation UMD-DYSF) (Krahn *et al*, 2009), without any apparent mutational hotspots. At least, a third of these mutations are missense mutations (Cacciottolo *et al*, 2011; Jin *et al*, 2016; Schoewel *et al*, 2012), leading for some to the production of misfolded proteins, aggregation (Spuler *et al*, 2008; Wenzel *et al*, 2006) and subsequent degradation by the ER-associated protein degradation machinery through the ubiquitin-proteasome system (ERAD I) or the alternative autophagy/lysosome degradation machinery (ERAD II) (Fujita *et al*, 2007). Although no treatment is currently available for LGMD R2, research directed towards missense protein refolding, or targeting cellular quality control to inhibit premature degradation of proteins in the ER with small molecules, opens up a therapeutic avenue for LGMD R2. Evidence in support of this strategy for rescuing missense mutations that cause LGMD has recently been described by our group for LGMD R3, as revealed by the positive impact of thiostrepton(Hoch *et al*, 2019) and the combination of givinostat and bortezomib on α-sarcoglycan missense mutant protein reallocation to the plasma membrane (Hoch *et al*, 2022), by disrupting proteasomal and autophagic degradation systems. Among the large panel of missense *DYSF* mutations that cause LGMD R2, we focused on the well characterized *DYSF*^L1341P^ mutant for our proof of concept. This mutant was identified 15 years ago (Wenzel *et al*, 2006) and initially shown to cause patchy sarcolemmal immunostaining and intracellular aggregates of dysferlin in the muscle fibers of patients. In 2007, Fujita et al. reported that aggregates of this dysferlin mutant on the ER membrane stimulated autophagosome formation via the activation of ER stress-eIF2α phosphorylation (Fujita *et al*, 2007). These results subsequently led to the investigation of pharmacological compounds that modulate autophagy, on one hand showing that the autophagy inducer rapamycin inhibited the accumulation of the *DYSF*^L1341P^ mutant, reducing its aggregation from 38% to 18%. On the other hand, the autophagy inhibitor 3-methyladenine (3-MA) was shown to increase accumulation from 36% to 44%. Later relocating this mutant form of dysferlin in the membrane with peptides was reported to restore part of its functionality, including the defective membrane repair capacities of muscular cells (Schoewel *et al*, 2012). More recently, based on the hypothesis that rescuing mutant forms of dysferlin could be a valuable therapeutic option, another study evaluated a selection of compounds using a 96-well plate cell-based flow cytometry assay that quantifies dysferlin localization in the plasma membrane (Tominaga *et al*, 2022). In this study, the authors highlighted the positive effect of the CFTR corrector corr-2b and the chemical chaperone 4-phenylbutyric acid (4-PBA) in rescuing *DYSF*^W992R^, *DYSF*^E1335G^, *DYSF*^L1341P^, and *DYSF*^F1867L^ mutants’ localization in the plasma membrane and, more particularly, the functional benefit of 4-PBA treatment in restoring the membrane repair capacities of myotubes expressing the L1341P dysferlin.

We report here the development of a high-throughput assay to identify any drugs capable of reallocating the aggregated *DYSF*^L1341P^ mutant form of dysferlin, by hypothesizing that this treatment might secondarily improve membrane repair function in patients’ cells. Taking advantage of a *DYSF*^L1341P^ immortalized myoblast cellular model, recapitulating aggregation of dysferlin in ER and the pathological phenotype of disrupted membrane repair (Philippi *et al*, 2012), we screened a library of 2239 drug candidates comprising off-patent small molecules, of which 95% are approved drugs, and bioactive compounds. Following primary screening and secondary assays, our results describe the discovery of the positive impact of two compounds, saracatinib and bazedoxifene, on reallocating misfolded L1341P dysferlin outside the ER and improving myoblasts resistance to osmotic shock. Finally, functional e*x vivo* analysis also revealed an additional protective effect of bazedoxifene on dysferlin-deficient myofibers from Bla/J mice carrying a KO mutation, opening a new avenue toward genotype-independent treatment of LGMD R2 patients.

## RESULTS

### Development and optimization of a high-content screening assay

A 384-multiwell screening assay was developed on LGMD R2 immortalized myoblasts to identify pharmacological compounds capable of rescuing L1341P dysferlin aggregation and reallocating the protein to the plasma membrane. The assay was based on the detection of dysferlin-positive cells by immunostaining and their quantification by high-content imaging, as described in Figure 1A. Briefly, immunofluorescence was performed using the desmin and dysferlin antibodies and images were acquired and analyzed using the CellInsight CX7 HCS automate and the HCS Studio software suite. An analysis algorithm based on a Colocalization Bioapplication was used to quantify dysferlin-positive cells, i.e. the number of myoblasts containing more than one area of dysferlin aggregates in their cytoplasm. Based on the intensity level of fluorescent signals, detection masks were defined to delimit the nuclei (Hoechst), to segment the myoblasts (Desmin) and to identify areas of dysferlin aggregates (Dysferlin) (Figures 1B and S1A). Dysferlin intensity detection threshold was chosen, so that less than 10% of cells were dysferlin-positive for the negative control (0.1% DMSO). As assessment of the robustness of the screening procedure requires identification of a positive control, several drugs targeting proteostasis were evaluated. To do so, the effect of proteasome inhibitors (Figure S1B), ERAD inhibitors (Figure S1C) and autophagy flux modulators (Figure S1D) was measured on *DYSF*^L1341P^ myoblasts, showing a significant increase of up to 60% in the number of dysferlin-positive cells after 24 hours of treatment with bafilomycin and chloroquine. Dose-response experiments were performed using these two autophagy inhibitors, bafilomycin and chloroquine, confirming a dose-dependent effect of these drugs on the number of L1341P dysferlin-positive myoblasts. The most efficient treatment was bafilomycin, inducing an increase in dysferlin-positive cells with an EC_50_ of 20nM and reaching maximum efficacy without toxicity at 64nM (Figure 1C). Chloroquine was also a potent drug but exhibited more toxic side effects on *DYSF*^L1341P^ myoblasts (data not shown). At last, quantification of dysferlin-positive cells in *DYSF*^L1341P^ myoblasts treated with 0.1% DMSO or increasing concentrations of bafilomycin was carried out to assess the robustness of the assay through calculation of a Z’ factor. The concentrations of bafilomycin that were compatible with high-throughput screening were the two most effective doses of 64 and 160nM, leading to Z’ factors of 0.58 and 0.63, respectively (Figure 1D).

**Figure 1.**
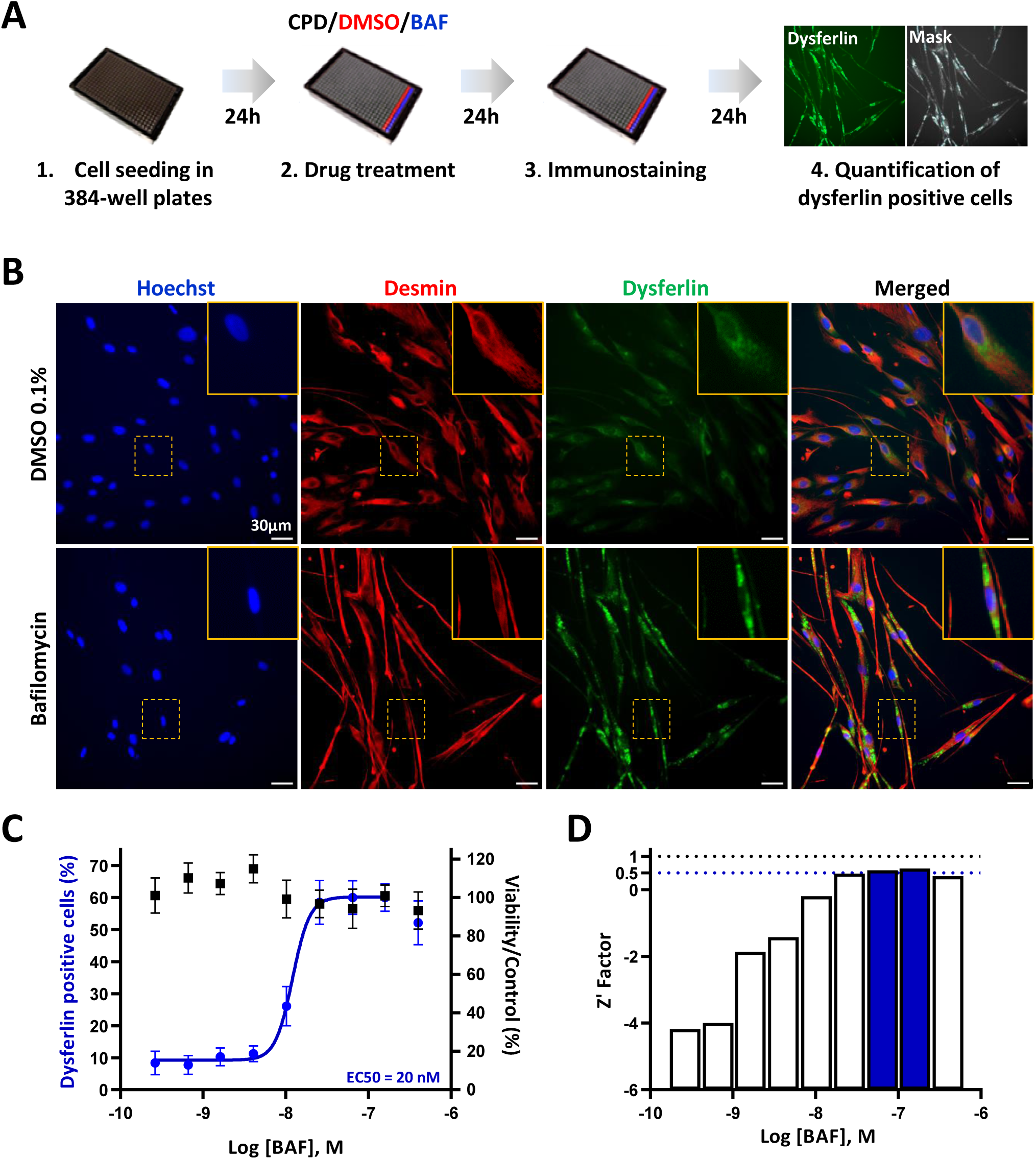
Development of a high-content screening assay to quantify the effect of drugs on *DYSF*^L1341P^ mutant expression. (**A**) Workflow for the high-content screening of L1341P dysferlin expression in 384-well plates. (**B**) Representative images of desmin (red) and dysferlin (green) staining in *DYSF*^L1341P^ immortalized myoblasts treated with 0.1% DMSO or 64 nM bafilomycin. Hoechst staining (blue) labels nuclei. Yellow outsets represent higher magnification images. Scale bar = 30 µm. (**C**) Quantification of dysferlin-positive myoblasts (blue) and cell viability (black) following treatment with increasing concentrations of bafilomycin. Each point represents the mean ± SD (n=8) of a representative experiment over four independent experiments and viability is expressed as percentage of the response induced by 0.1% DMSO. Half-maximal effective concentration (EC50) is indicated. (**D**) Determination of the Z′ factor for each concentration of bafilomycin tested. Blue bars represent the concentrations required to obtain a Z′ factor ≥ 0.5. Abbreviations: CPD, tested compounds; BAF, bafilomycin.

### Identification of drug candidates for rescuing misfolded L1341P dysferlin from degradation using high-throughput screening

A total of 2239 compounds were screened at 5 μM for their capacity to rescue L1341P misfolded dysferlin from degradation in *DYSF*^L1341P^ myoblasts. This library comprises 1280 FDA- and EMA-approved drugs from the Prestwick Chemical library and 959 FDA-approved drugs and bioactive compounds from the Selleckchem library. Quality of the screening assay was ensured by a statistically relevant difference between the negative 0.1% DMSO and positive 64nM bafilomycin controls (Figure 2A) that was measured for each of the 8 plates, giving a mean Z’ factor of 0.55 (Figure 2B). Compounds were considered as potential candidates when their effect was superior to five standard deviations from the mean for all the tested compounds, without affecting cell viability by more than 50% (Figure 2C). This led to an initial list of 13 hits (Figure 2D), with the number of L1341P dysferlin-positive myoblasts comprising between 24% and 53% for the least and most effective compound, respectively. Successive retest experiments were then performed to validate the effect of these 13 candidates on the dysferlin read-out (Figure 3A). A first retest step was conducted in quadruplicate on *DYSF*^L1341P^ myoblasts under the same screening conditions for each of the 13 compounds, identifying PHA-665752 and SRT1720 as false positives with less than 24% of dysferlin-positive cells after 24 hours of treatment (Figure 3B). Moreover, detection of green-stained myoblasts without the dysferlin primary antibody after 24 hours of treatment with obatoclax mesylate revealed fluorescence interference for this drug and thus excluded it from the list of primary compounds (Figure S2). Dose-response experiments were then carried out on the remaining 10 confirmed hits, with increasing concentrations from the nanomolar to micromolar range (Figure 3C). Of these, the drug quinacrine was discarded because of the absence of efficacy on the dysferlin-positive cell readout. The 9 remaining compounds were validated as hits of interest with an EC_50_ comprised between 345nM to 3.9µM. Of these, GSK1070916, NVP-BSK805, saracatinib and bazedoxifene induced a maximal effect equal to or greater than 60% on dysferlin-positive cell numbers after 24 hours of treatment with their maximum effective dose, 2µM or 5µM, and an associated toxicity of less than 20%. Crenolanib and SGI-1776 were the least effective compounds, with 24% and 30% dysferlin-positive cells, respectively, and a toxicity of 40% for crenolanib.

**Figure 2.**
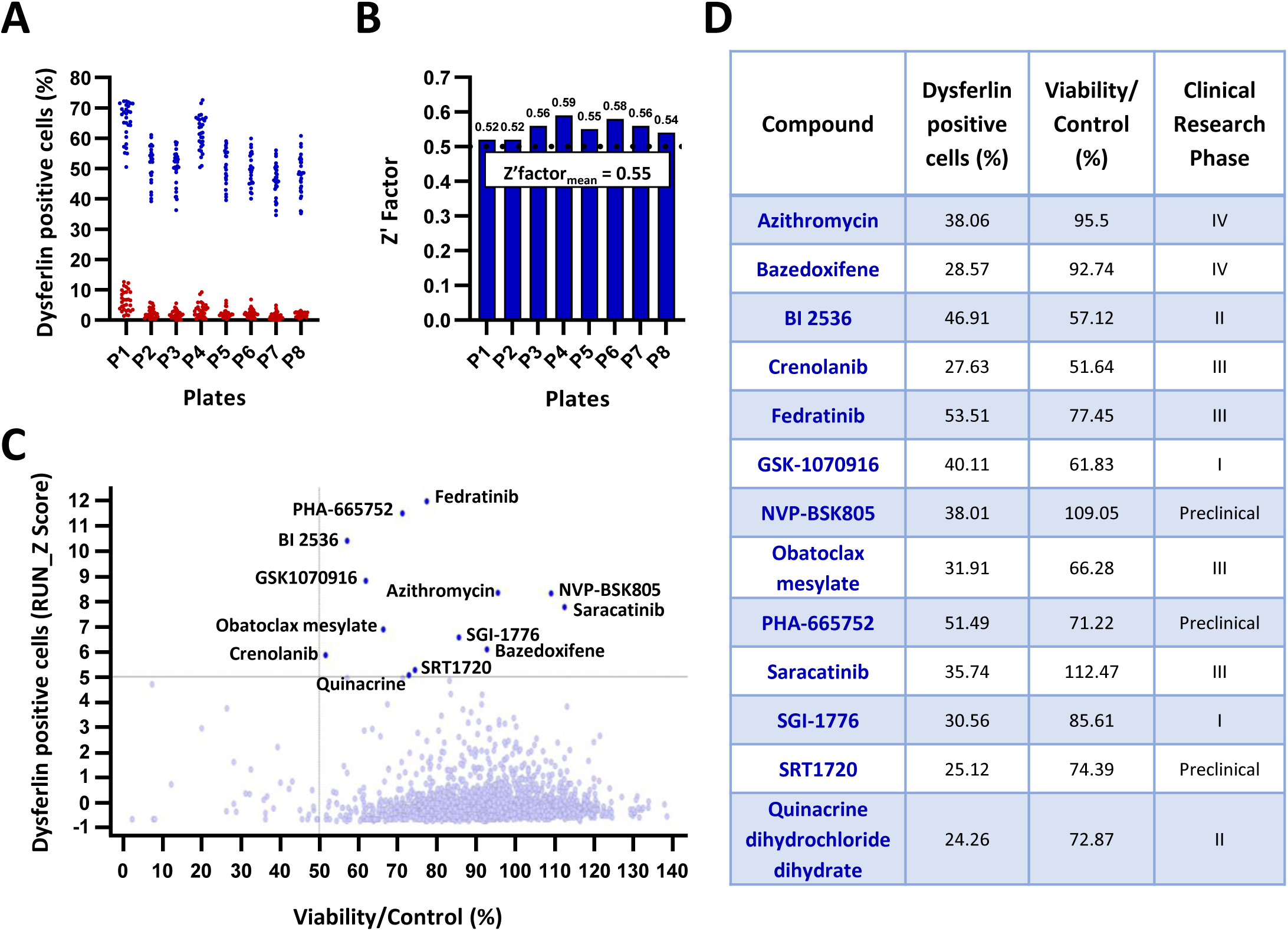
High-content screening for L1341P dysferlin expression. (**A**) High-throughput screening validation of L1341P dysferlin expression in myoblasts treated with the negative control, 0.1% DMSO (red), or the positive control, 64 nM bafilomycin (blue), in each of the 384-well plates used for screening (32 replicates/plate). (**B**) Determination of the Z′ factor for each of the 384-well plates used for screening and measurement of the mean for all plates. Blue bars represent screening plates with a Z′ factor ≥ 0.5. (**C**) Primary screen cell-based assay for L1341P dysferlin expression. Dot plot representation of the effects of the 2239 drugs on the number of dysferlin-positive cells (Z-score>5) and cell viability (Viability >50%). (**D**) List of the 13 compounds identified as primary hits during screening and the corresponding percentage of dysferlin-positive cells, percentage of viability and the maximum clinical research phase achieved by the compound.

**Figure 3.**
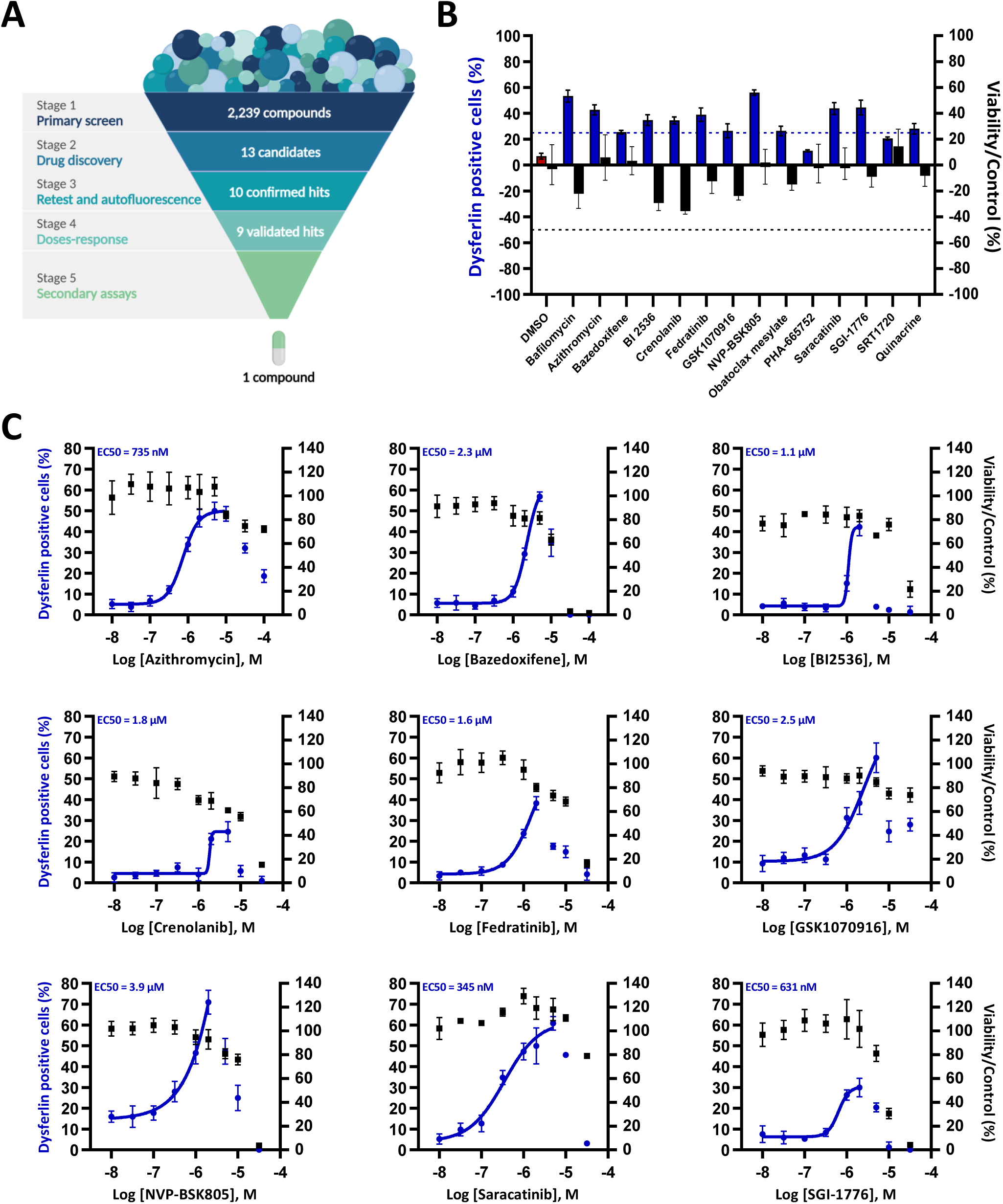
Validation of the primary hits on L1341P dysferlin expression. (**A**) Schematic representation of the retest experiments to validate the effect of the 13 primary candidates that were identified. (**B**) Hit confirmation: quantification of dysferlin-positive cells (blue) and cell viability (black) following 24 hours of treatment with the candidate compounds at the single 5µM dose used in screening. Each bar represents the mean ± SD (n=4) of a representative experiment over three independent experiments. (**C**) Hit validation: quantification of dysferlin-positive cells (blue) and cell viability (black) following treatment for 24 hours with increasing concentrations of confirmed hits. Each graph is a representative experiment (n=3). Each point represents the mean ± SD of four replicates. Half-maximal effective concentration (EC50) is indicated for each dose-response curve, modeled using nonlinear regression with 95% confidence level.

### Effect of validated hits on *DYSF*^L1341P^ mutant expression and localization

To determine whether the effect of the 9 validated compounds on the *DYSF*^L1341P^ mutant protein is transcriptional or post-transcriptional in origin, the *DYSF* gene expression was evaluated on L1341P myoblasts after 24 hours of treatment with the maximum effective dose of the drugs, as determined previously by dose-response experiments (10µM: azithromycin; 5µM: bazedoxifene, crenolanib, GSK1070916, saracatinib; 2µM: BI2536, fedratinib, NVP-BSK805, SGI-1776) (Figure S3). Analysis of qPCR revealed no significant difference following treatment with azithromycin, bazedoxifene, crenolanib, GSK1070916, NVP-BSK805, saracatinib and SGI-1776 in comparison to 0.1% DMSO. In contrast, *DYSF* gene expression was significantly reduced after treatment with BI2536 (∼0.5-fold change) or increased after treatment with fedratinib (∼1.7-fold change), indicating a slight transcriptional deregulation in the *DYSF* gene. Dysferlin protein level and subcellular localization were also investigated by immunostaining after treatment with the 9 validated hits, at the same concentrations as those used for gene expression analysis (Figure 4). Endoplasmic reticulum was labeled using the KDEL antibody to monitor the capacity of the drugs to relocate L1341P dysferlin outside this organelle. Confocal analysis revealed that treatment of *DYSF*^L1341P^ myoblasts with each of the 9 drugs leads to an increase in dysferlin expression and to a relocalisation of staining outside of the endoplasmic reticulum.

**Figure 4.**
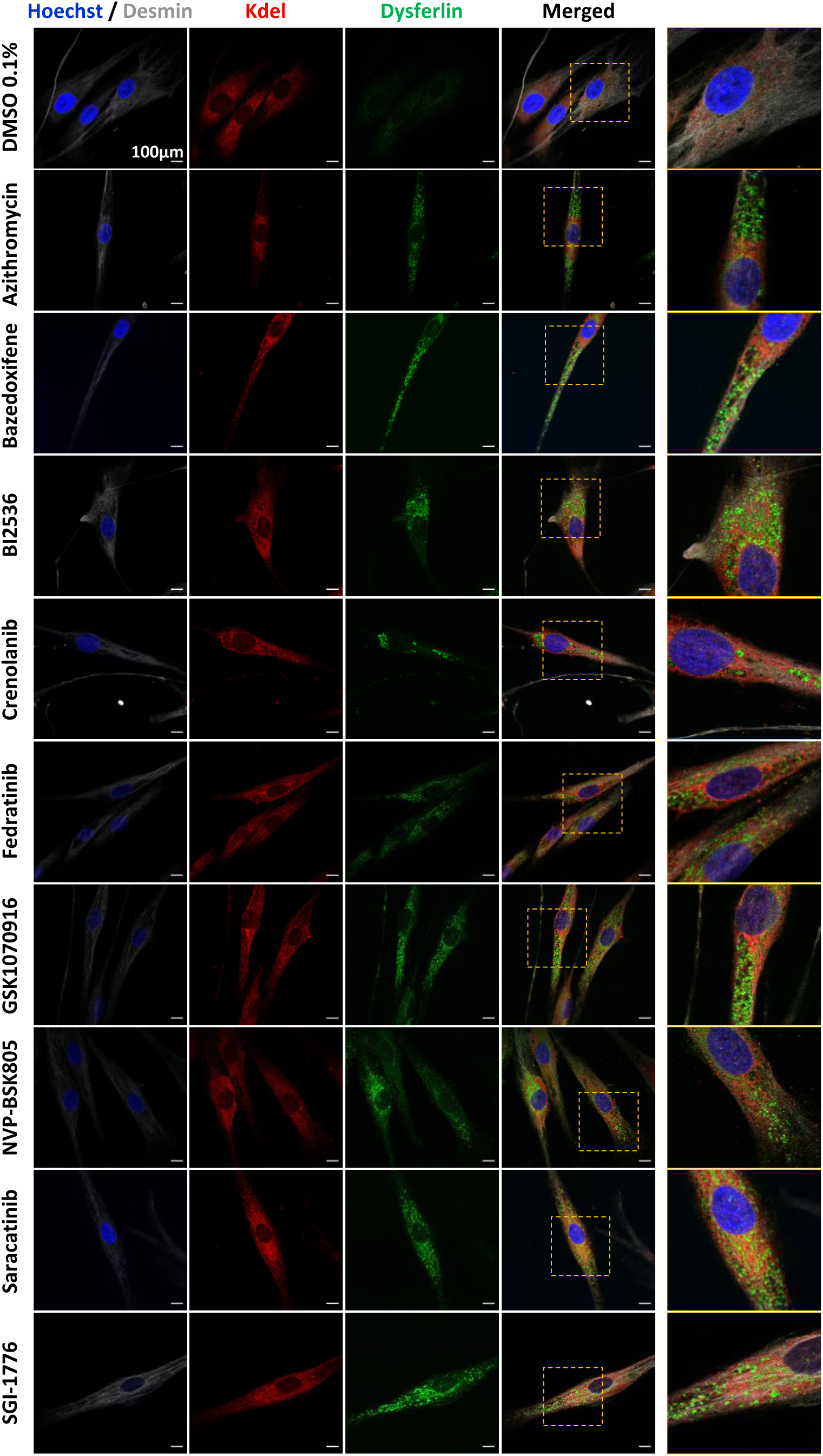
Evaluation of the effect of validated hits on L1341P dysferlin protein localization. Representative images of desmin (white), KDEL (red) and dysferlin (green) immunostaining in *DYSF*^L1341P^ myoblasts treated for 24 hours with the control 0.1% DMSO or the 9 validated compounds at the dose of 5 µM. Nuclei are labeled by Hoechst staining (blue). Yellow outsets represent higher magnification images. Scale bar = 100 µm.

### Identification of two molecules, saracatinib and bazedoxifene, that improve the impaired membrane repair phenotype

The major pathological feature in LGMD R2 is compromised membrane repair. To explore the potential of identified compounds to correct the dysferlin functional defect, we thus measured the ability of cells to repair their plasma membrane following stress by performing a hypo-osmotic shock assay (Barthélémy *et al*, 2015; Kerr *et al*, 2013). First, osmotic shock assay was developed by adding a hypotonic solution to healthy human myoblasts that had been previously transfected with a dysferlin-specific small interfering RNA (siRNA), inducing a reduction in protein compared to the scrambled siRNA negative control (Figure S4A). Plasma membrane integrity was monitored by following cell mortality (i.e. the number of cells positive for green dye, a marker of apoptosis) over time compared to the number of cells at the beginning of experiment (i.e. the number of cells positive for red dye, a nuclear marker) (Figure S4B). The percentage of cell mortality in healthy myoblasts that had been pre-treated with scrambled siRNA (or not) increased from 50% after 12 hours of osmotic shock to 100% after 16 hours (Figures S4B and S4C), confirming osmotic shock-induced mechanical stress on the cell membrane, as previously reported (Barthélémy *et al*, 2015; Pajovic *et al*, 2016; Barzilai-Tutsch *et al*, 2018). Conversely, osmotic shock on myoblasts lacking dysferlin induced an exacerbated mortality that reached 50% after 6 hours and 100% after 12 hours (Figures S4B and S4C), demonstrating the link between dysferlin expression and its protective role in the plasma membrane against osmotic shock. Using this procedure, functionality of the 9 validated hits was thus assessed in *DYSF*^L1341P^ myoblasts pre-treated with the compounds at EC_50_ for 24 hours (Figure 5A). Of these molecules, our results indicated that saracatinib and bazedoxifene improved the resistance of cells to osmotic shock, with 50% mortality in 20 hours and more than 50 hours, respectively, compared to 15 hours after treatment with 0.1% DMSO (Figure 5B and 5C). The effect of increasing concentrations of these two molecules, saracatinib and bazedoxifene, was evaluated in *DYSF*^L1341P^ myoblasts, demonstrating the dose-response effect of the compounds on the cells’ resistance to osmotic shock (Figure 5D and 5E). At 2µM and 1µM, the maximum effective dose for saracatinib and bazedoxifene, osmotic shock induced 50% cell mortality in 37 hours and 63 hours, respectively, compared to about 10 hours with the 0.1% DMSO control. The specificity of the effect of saracatinib and bazedoxifene on osmotic shock resistance was then assessed in healthy (Figure 5F), L1341P dysferlin (Figure 5G) and null-dysferlin (Figure 5H) immortalized myoblasts that had been pre-treated with the maximum effective dose of the two compounds. At the 2µM concentration, our results described that saracatinib improved survival of L1341P dysferlin myoblasts but not that of healthy or null-dysferlin myoblasts following osmotic shock, while bazedoxifene (1µM) was efficient in all three models. Thus, these observations suggest that the effect of saracatinib on cell resistance to osmotic shock would be related to the rescue of the misfolded L1341P dysferlin, whereas bazedoxifene could have an additional and unrelated positive effect on cell survival.

**Figure 5.**
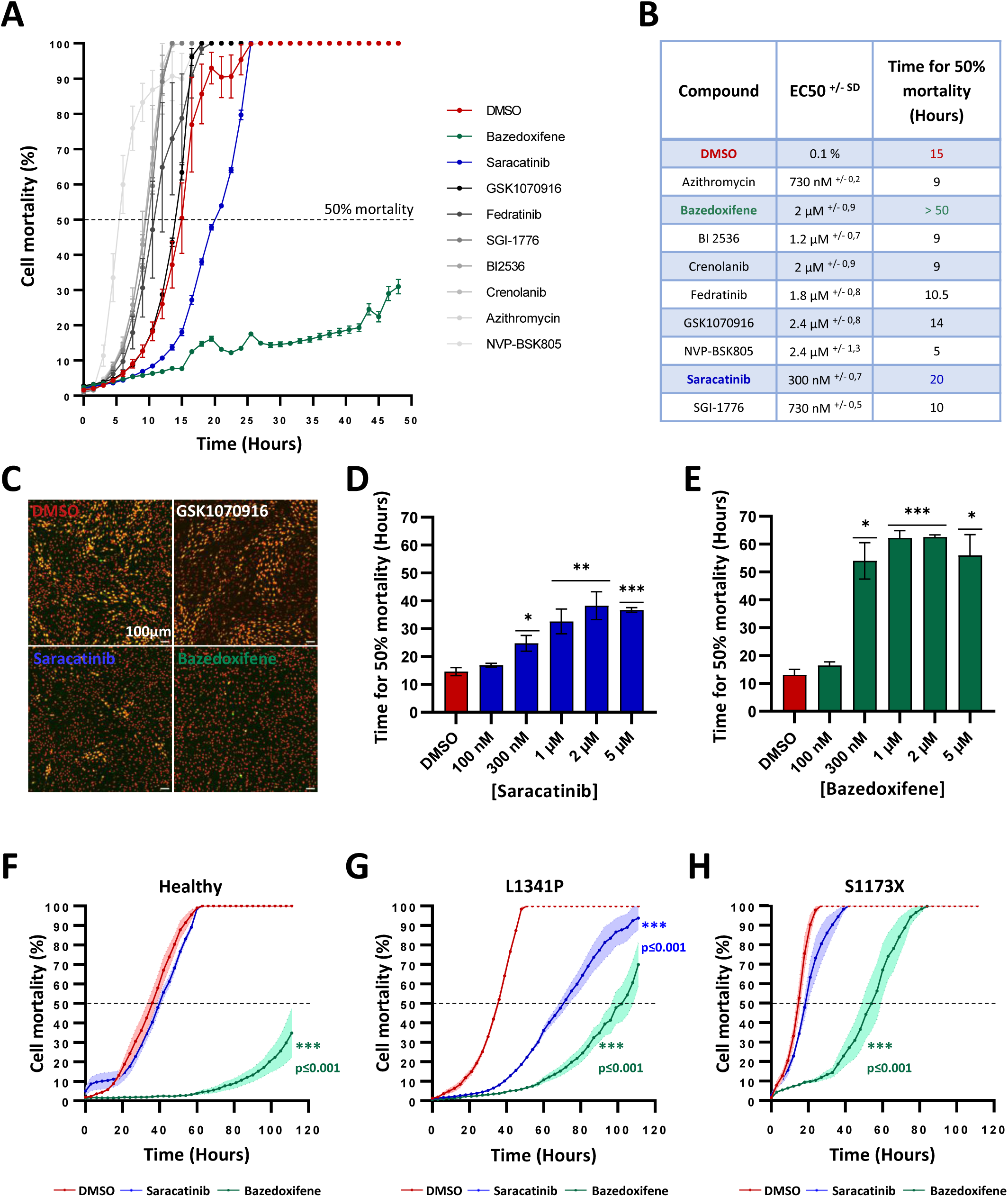
Efficacy of compounds on the *DYSF*^L1341P^ impaired membrane repair phenotype. (**A**) Measure of cell mortality following hypo-osmotic shock on *DYSF*^L1341P^ myoblasts pre-treated for 24 hours with the 9 validated compounds at their half-maximal effective concentration (EC_50_). Each point represents the mean ± SD (n=4) of a representative experiment over three independent experiments. (**B**) Table of the osmotic shock duration required to reach 50% mortality for each of the 9 compounds. (**C**) Representative images of automated quantification of cell mortality (green/red dyes ratio) after 15 hours induction of osmotic shock on *DYSF*^L1341P^ myoblasts pre-treated with molecules, when 50% mortality is achieved with 0.1% DMSO. Scale bar = 100 µm. (**D-E**) Dose-response experiments. Measure of time required to reach 50% mortality following hypo-osmotic shock on *DYSF*^L1341P^ myoblasts pre-treated for 24 hours with increasing concentrations of saracatinib (**D**) or bazedoxifene (**E**). Each chart represents the mean ± SD (n=4) of a representative experiment. \**p* ≤ 0.05, \*\**p* ≤ 0.01, *** *p* ≤ 0.001 (Brown-Forsythe and Welch ANOVA with Dunnett’s T3 multiple comparisons test). (**F-H**) Quantification of cell mortality following hypo-osmotic shock on healthy (**F**), *DYSF*^L1341P^ (**G**), or *DYSF*^S1173X^ (**H**) immortalized myoblasts pre-treated for 24 hours with 0.1% DMSO (red curve) or the maximum effective dose of saracatinib (2µM, blue curve) or bazedoxifene (1µM, green curve). Each point represents the mean ± SD (n=4) of a representative experiment. *** *p* ≤ 0.001 (One-way ANOVA with Dunnett’s multiple comparisons test).

### Bazedoxifene treatment improves muscle fibers resistance of dysferlin KO mice

Evaluation of membrane repair was then performed on myofibers isolated from flexor digitorum brevis (FDB) muscles of dysferlin KO (Bla/J) and control (C57BL/6J) male mice. Osmotic shock assay was induced in the presence of FM1-43 dye which enters the myofiber upon membrane disruption, causing an increase in fluorescence intensity as FM1-43 binds to the internal membrane(Defour *et al*, 2014). Plasma membrane integrity can thus be monitored for each fiber across all the timepoints of the experiment through measurement of fluorescence intensity. Indeed, in the absence of any treatment, a higher fluorescence intensity over time was observed in Bla/J fibers compared to those isolated from control mice, confirming worse repair ability in dysferlin-null muscle fibers (Figure S5). To evaluate whether drug treatment is beneficial to improve membrane repair capacity, Bla/J and control myofibers were treated with bazedoxifene (1µM) or PBS for 40 minutes prior to osmotic shock assay. While no significant differences in FM1-43 fluorescence intensity was observed in control myofibers treated with bazedoxifene compared to PBS, a decrease dye entry was monitored in those isolated from Bla/J mice (Figure 6A), highlighting an improvement of plasma membrane repair with bazedoxifene treatment. These results were confirmed by calculating the coefficient of saturation, which reflects the fluorescence intensity per time unit, for each myofiber in each animal. This coefficient was not modulated in control myofibers treated with bazedoxifene compared to PBS, with mean values of 22.8 and 22.2 ΔF A.U./min, respectively (Figure 6B). Conversely, coefficient of saturation significantly decreased in Bla/J fibers from 25 following PBS treatment to 15.33 with bazedoxifene (Figure 6C). All together, these data suggest a protective role of bazedoxifene against membrane damage in dysferlin-null muscle fibers.

**Figure 6.**
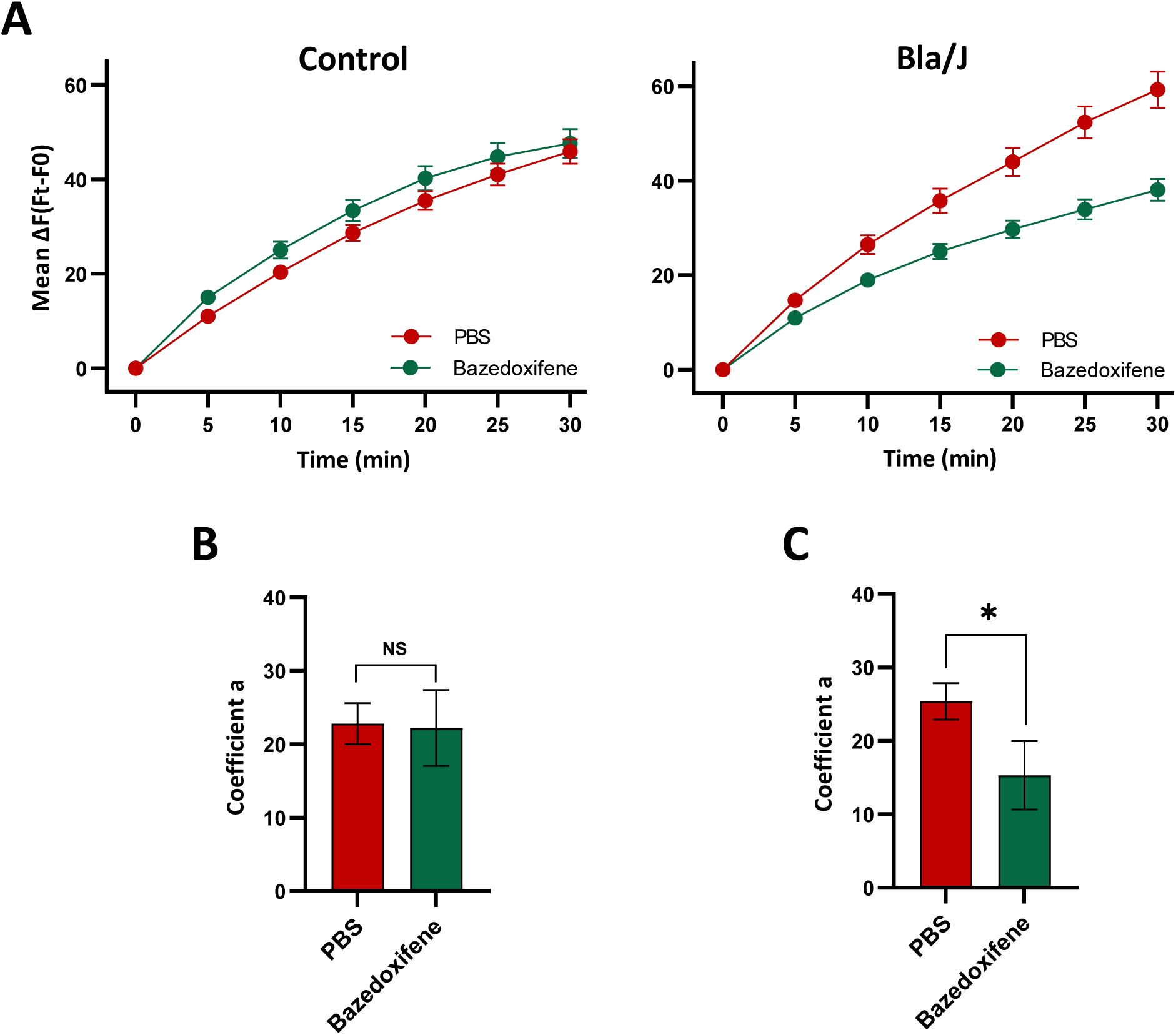
Treatment with bazedoxifene improves membrane repair capacity of isolated Bla/J muscle fibers. (**A**) Quantification of intracellular FM1-43 dye fluorescent intensity in PBS- (red) or 1µM bazedoxifene- (green) treated FDB fibers isolated from control (left panel) and Bla/J (right panel) animals after osmotic shock induction. Values are measured as the difference in fluorescence intensity between t and t0 (ΔF = Ft-F0). The data are represented as mean ΔF ± SEM of fibers isolated from 6 animals per group (126 ≤ fibers ≤ 287). (**B-C**) The data were fit to a model as ΔF= 1+a log(tn+1)+b. The coefficient of saturation “a” is the reflection of the increase of fluorescence intensity per time unit in control (**B**) and Bla/J (**C**) myofibers, treated or not with bazedoxifene. * *p* ≤0.05 (Student’s t-test).

### Bazedoxifene functions as an autophagy flux inducer

Bazedoxifene is a third-generation selective estrogen receptor modulator (SERM) approved for the prevention of postmenopausal osteoporosis. Because Bazedoxifene and SERMs were previously described as potent inducer of autophagy (Torres-López *et al*, 2019; Kim *et al*, 2015; Xiang *et al*, 2019) and given that autophagy plays a key role on both proteostasis and membrane repair, we hypothesized that bazedoxifene’s effect on LGMD R2 was triggered by autophagy induction. To investigate this potential mechanism of action, we first quantify autophagic vacuoles in live cells using the Cyto-ID® Autophagy detection kit. Treatment with PP242, a known autophagy inducer, was used as a positive control and compared to the negative DMSO 0.1% control showing an increase of dye staining response in myoblasts (Figure S6A). The autophagic signal, calculated with median fluorescence intensity (MFI), was 2.0 and 1.5 times greater in healthy and *DYSF*^S1173X^ myoblasts treated with PP242 (Figure S6B), respectively. With this method, we then demonstrated that bazedoxifene treatment induced an accumulation of autophagic vesicles in myoblasts (Figure 7A), with a threefold increase in MFI compared to the control (Figure 7B). To confirm these observations, we also measured by immunostaining the effect of bazedoxifene on LC3B, a protein known to be in stringent association with the autophagosomal membranes formed during the autophagic process (Runwal *et al*, 2019). As a control and to validate our quantification method, we used chloroquine, a late autophagy inhibitor that prevent the activity of lysosomal acid proteases and cause autophagosomes to accumulate (Mauthe *et al*, 2018) (Figure S6C). As shown, increasing concentrations of chloroquine led to dose-dependent effect on LC3B fluorescent signal (Figure S6D and E). Using this method, we then demonstrated that bazedoxifene treatment induced the appearance of LC3B-positive dots compared to the basal DMSO 0.1% condition (Figure 7C), as well as a significant 15- to 30-fold increase in total LC3B puncta area (Figure 7D) and 10- to 20-fold increase in LC3B puncta intensity (Figure 7E), in healthy and *DYSF*^S1173X^ myoblasts, respectively. To further characterize the effect of bazedoxifene on autophagy, we then monitored LC3B aggregation in autophagosomes and autolysosomes using a RFP-GFP-LC3B tandem sensor. This sensor combine an acid-sensitive green fluorescent protein (GFP)-LC3B reporter with an acid-insensitive red fluorescent protein (RFP)-LC3B reporter, allowing to visualize the transition from neutral autophagosomes to acidic autolysosomes. Accordingly, healthy and *DYSF*^S1173X^ patient’s myoblasts were transiently transduced with the RFP-GFP-LC3B tandem sensor and treated with 5µM bazedoxifene or DMSO 0.1%. Our results revealed that bazedoxifene treatment significantly increased the number of GFP- and RFP-LC3B structures (Figure 7F), reflecting the accumulation of acid and non-acid autophagic structures. Based on these data, we then evaluated the conversion of LC3B from the soluble cytosolic form, LC3-I, to the lipidated membrane-associated form, LC3-II (Kabeya *et al*, 2000), using immunoblot analysis. Our results revealed that bazedoxifene treatment induced the conversion to LC3-II in healthy and *DYS*F^S1173X^ myoblasts (Figures 7G and 7H), reflecting an accumulation of autophagosomes. However, autophagic compartments are intermediate constituents of a dynamic lysosomal degradation process and their intracellular abundance detected at a given time point is a function of the balance between the rate of their generation and the rate of degradation. Accordingly, increased LC3-II could reflect either increased autophagosome formation due to early autophagy induction or a blockage in the downstream steps in autophagy (Rubinsztein *et al*, 2009). Thus, to confirm that bazedoxifene is an autophagy inducer, LC3-II turnover was then analyzed in the presence and absence of ammonium chloride (NH4Cl), an autophagy inhibitor that raises the luminal pH of intracellular vesicles and prevents the activation of degradative enzymes inside lysosomes (Klionsky *et al*, 2012). As hypothesized, immunoblot analysis revealed a higher increase of LC3-II levels in both healthy and *DYSF*^S1173X^ treated with the combination of bazedoxifene and NH4Cl (Figure 7G), compared with compounds alone, indicating that the increased accumulation of LC3-II induced by bazedoxifene is not due to an inhibition of downstream autophagic flux. To confirm that bazedoxifene induces autophagy, treatment with another inhibitor, 3-methyladenine (3’MA), was used to block upstream autophagy. As expected, immunoblot analysis revealed that treatment with 3’MA partially reverse the effect of bazedoxifene on LC3-II conversion in both healthy and *DYSF*^S1173X^ myoblasts (Figure 7H).

**Figure 7.**
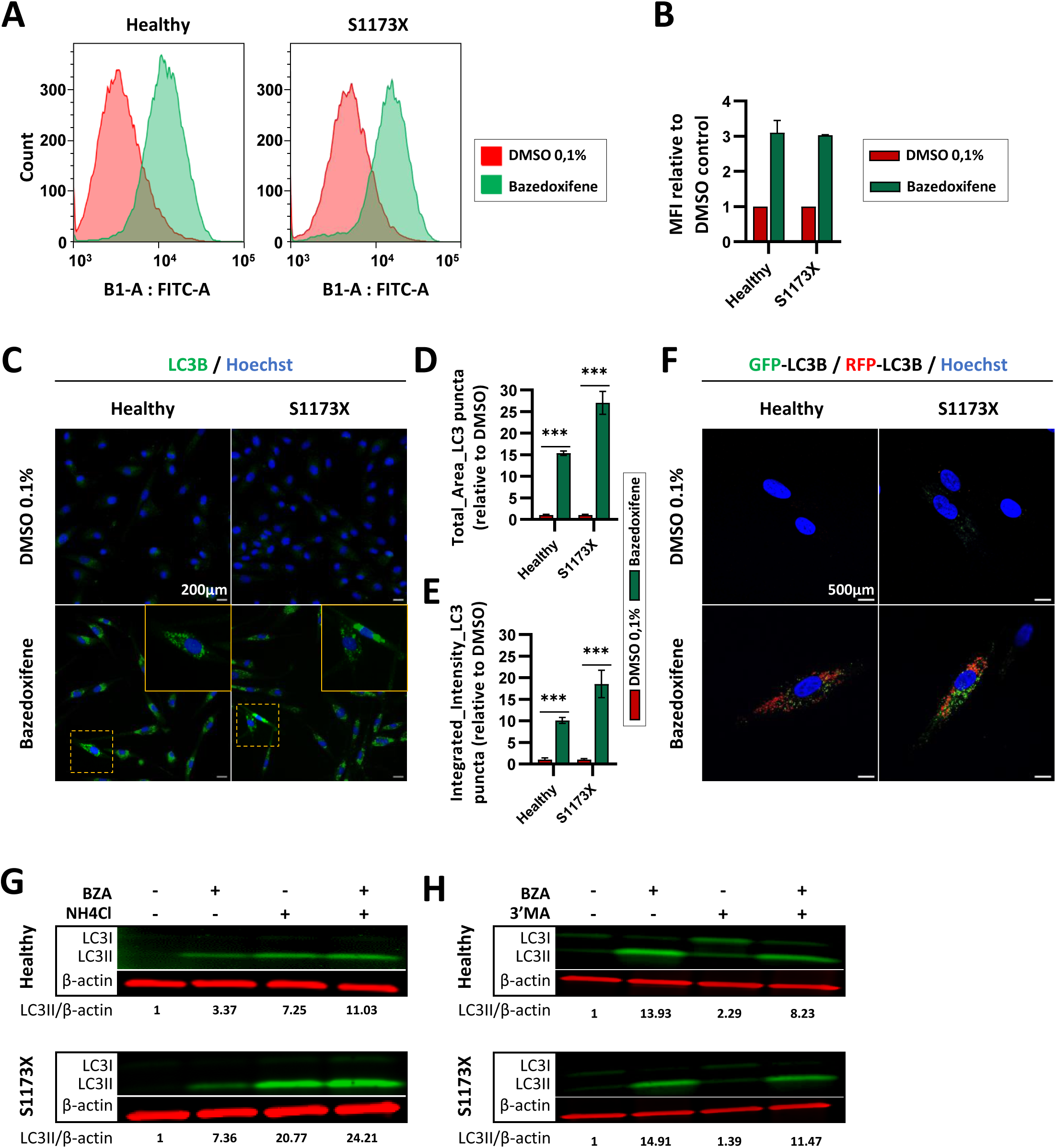
Characterization of autophagy flux in LGMD R2 myoblasts treated with bazedoxifene. (**A-B**) Measure of autophagy flux by flow cytometry. Representative FACS plots showing eGFP fluorescence detected using a CYTO-ID® Autophagy detection kit in healthy and *DYSF*^S1173X^ myoblasts, treated with or without bazedoxifene (5µM) for 24 hours (**A**). Median fluorescence intensity (MFI) was assessed for each condition and normalized to DMSO control (**B**). Data are shown as the mean ± SD (n=2). (**C-E**) Formation of LC3 positive structures under control of bazedoxifene treatment. Representative immunofluorescence images showing LC3B staining (green) in healthy and *DYSF*^S1173X^ immortalized myoblasts treated with 0.1% DMSO or 5µM bazedoxifene for 24 hours (**C**). Hoechst staining (blue) labels nuclei. Yellow outsets represent higher magnification images. Scale bar = 200 µm. Quantification of the relative LC3-positive structures’ average total area (**D**) and integrated intensity (**E**), using MetaXpress software. Each bar represents the mean ± SD (n=4 or more) of a representative experiment over three independent experiments. \*\*\**p*≤0.001 (2way ANOVA multiple comparisons). (**F**) Representative confocal images depicting GFP-LC3 (green) and RFP-LC3 (red) puncta in myoblasts transiently expressing GFP-RFP-LC3B and treated with bazedoxifene (5µM) or DMSO (0.1%) for 24 hours. Nuclei are labelled by Hoechst staining (blue). Scale bar = 500µm. (**G-H**) Representative immunoblot analysis of LC3I and LC3II expression in healthy and *DYSF*^S1173X^ myoblasts treated with DMSO (0.1%) or bazedoxifene (5µM) for 24 hours, in absence or in presence of 5mM NH4Cl for 24 hours (**G**) or 3mM 3’MA for 2 hours (**H**). The values below the blot show relative levels of LC3II/β-actin protein after normalization to DMSO-treated cells. Abbreviations: BZA, bazedoxifene; 3’MA, 3-Methyladenine.

## DISCUSSION

The main result of this study is the identification of the protective effect of two drugs, saracatinib and bazedoxifene, as potential therapeutic options for the treatment of LGMD R2 patients carrying the *DYSF*^L1341P^ mutation. While our drug screening described a positive effect of nine drugs on L1341P dysferlin aggregation, secondary assays highlighted the therapeutic potential of two drugs in also rescuing osmotic shock resistance in LGMD R2 cells. As previously described, the capacity of LGMD R2 cells to resist osmotic shock reflects their membrane repair properties (Kerr *et al*, 2013; Barthélémy *et al*, 2015). We report here that the effect of saracatinib on osmotic shock is only observed in cells expressing L1341P misfolded dysferlin, while bazedoxifene is efficient in all the cell lines, including healthy or dysferlin KO cells which suggests a distinct mechanism of action. Interestingly, these two drugs have been previously described to induce autophagy; an evolutionarily conserved catabolic process that participates in the degradation of damaged cellular organelles and misfolded protein aggregates in a cellular homeostasis context and in response to biological stress.

Developed by AstraZeneca as an anti-invasive and anti-tumor drug, saracatinib (AZD-0530) is a dual kinase inhibitor, with selective actions as a Src inhibitor and a Bcr-Abl tyrosine-kinase inhibitor. Because the inhibition of Src by siRNA or inhibitors promotes autophagy in cancer cells through Akt/mTOR signaling (Wu *et al*, 2010), several studies have investigated the use of saracatinib to clear aggregates as misfolded phosphorylated Tau protein (Nygaard *et al*, 2015; Tang *et al*, 2020) or α-synuclein aggregates (Choi *et al*, 2020) in Alzheimer’s and Parkinson’s diseases, respectively. This capacity to clear protein aggregates (dissolve and/or eliminate) might explain our data describing the positive impact of saracatinib on L1341P misfolded dysferlin and the absence of effect in healthy or dysferlin KO cells. Bazedoxifene is a third generation selective estrogen receptor modulator (SERM) (De Villiers *et al*, 2011), developed by Pfizer. In late 2013, Pfizer received approval for bazedoxifene as part of the combination drug DUAVEE for the prevention (not treatment) of postmenopausal osteoporosis. Thereafter, a few studies highlighted the potential of this drug to improve cell viability. In 2019, it was reported that bazedoxifene treatment prevented apoptosis of satellite cells through the estrogen receptor, suggesting a pro-survival effect on muscular cells (Collins *et al*, 2019). More recently, some studies suggested that bazedoxifene played a role as an inducer of autophagy, on one hand, through PDGF-BB regulation in vascular smooth muscle cells (Song *et al*, 2020) and, on the other hand, by inducing autophagosome formation and LC3B-II protein expression through Akt/mTOR signaling in macrophages infected with M. tuberculosis (Ouyang *et al*, 2020) or in polarized Th17 cells (Wang *et al*, 2020). Because bazedoxifene treatment stimulates LC3B vesicles production in LGMD R2 muscular cells and because LC3-II vesicles might be recruited to the damaged membrane site for membrane resealing process (Corkery *et al*, 2023), we hypothesized that its widespread protective effect might therefore be linked to this effect on autophagy rather than to the clearance of protein aggregates. This hypothesis is also supported by the fact that bazedoxifene belongs to the same class of drugs as tamoxifen, reported to improve muscle contractility in preclinical models of centronuclear myopathies (Gineste *et al*, 2023; Gayi *et al*, 2018b) and currently investigated in a phase 3 clinical trial to treat Duchenne muscular dystrophie (NCT03354039) (Gayi *et al*, 2018a; Dorchies *et al*, 2013; Nagy *et al*, 2019; Birnbaum *et al*, 2022; Kuşçu *et al*, 2022).

Since its discovery, autophagy has been shown to be involved in a wide range of pathological conditions, most notably in neurodegenerative diseases such as Alzheimer’s disease, amyotrophic lateral sclerosis, Parkinson’s disease, and Huntington’s disease. While displaying different symptoms, several studies have reported 1/ common autophagy impairments and 2/ a clear relationship between toxic protein aggregation and the progressive lysosomal dysfunction in these diseases (Nixon & Yang, 2011). This degradation process has been intensively studied in neurodegenerative diseases and plays a key role in maintaining cellular homeostasis in skeletal muscles, both during rest and exercise (Jokl & Blanco, 2016; Lira *et al*, 2013; Masiero *et al*, 2009). In myopathies, the main downstream effect of autophagy impairment is the accumulation of damaged cellular proteins and organelles that activate signaling cascades leading to fiber atrophy and apoptosis and, in some cases, to a reduction in the mechanical force generated during muscle contraction and a failure of muscle regeneration. Over the years, several treatments stimulating autophagy (such as starvation, low-protein diet, or pharmacologic inhibition of mTOR kinase) have been shown to have a positive impact in animal models of multiple myopathies, confirming that autophagy deficiency plays an important role in the pathogenesis of skeletal muscular disorders. Recently, promising results in a limited proof-of-principle clinical trial on a low-protein diet have been reported in eight adult patients with collagen VI–related congenital muscular dystrophies (NCT01438788). In addition to the intrinsic positive effect of autophagy inducers, our data suggest that saracatinib and bazedoxifene might target mechanistically distinct autophagy processes. While saracatinib may have a second positive impact in patients carrying the *DYSF*^L1341P^ mutation and other missense proteins degraded by the same pathway, bazedoxifene could be effective for a broader spectrum of pathologies associated with muscle cell fragility, particularly with regard to the integrity of their membranes.

## MATERIALS AND METHODS

### Myoblast cell lines and culture

Experiments were performed with human immortalized myoblasts from unaffected individuals or LGMD R2 patients. The LGMD R2 cell lines were established from muscle biopsies from patients bearing the relevant mutations: homozygous missense *DYSF* c.4022T>C; p.L1341P (*DYSF*^L1341P^) and heterozygous nonsense *DYSF* c.342-1G>A/c.3516_3517delTT; p.Ser1173X (*DYSF^S1173X^*). All cell lines were obtained from the immortalization platform of human cells from the Institut de Myologie (Paris, France), with agreement of the subjects through signature of an informed consent and anonymization before immortalization, according to the EU GDPR regulation. Immortalized myoblast cell lines were cultured in growth medium consisting of 1 volume of 199 medium (Invitrogen, United States) for 4 volumes of Dulbecco’s modified Eagle’s medium (Invitrogen, United States), supplemented with 20% fetal bovine serum (Sigma-Aldrich, United States), 25 µg/ml fetuin (Life Technologies, United States), 5 ng/ml epidermal growth factor (Life Technologies, United States), 0.5 ng/ml basic fibroblast growth factor (Life Technologies, United States), 0.2 µg/ml dexamethasone (Sigma-Aldrich, United States) and 5 µg/ml insulin (Sigma-Aldrich, United States). Cells were seeded on 0.1% gelatin-coated plates and maintained in a humidified atmosphere of 5% CO_2_ at 37°C until experimentation.

### Animal models

B6.A-Dysfprmd/GeneJ (strain: 012767; referred as Bla/J) and C57BL/6J (referred as WT or control) mice were housed in an SPF barrier facility with a 12h light/ dark cycle in the Center for Exploration and Experimental Functional Research, (CERFE, GIP GENOPOLE, France) and were provided with food and water ad libitum. All animals were handled according to French and European guidelines for human care and use of experimental animals.

### High-throughput screening

The high-throughput screening was conducted using a Biocel 1800 system (Agilent, United States). To this end, *DYSF*^L1341P^ myoblasts were plated in 38 μl of culture medium into black 384-well clear-bottom plates. 24 hours after seeding, 2 μl of 20x compounds from the chemical libraries were transferred into cell assay plates in monoplicate. In each plate, the negative control (0.1% DMSO) and positive control (64 nM bafilomycin) were added in columns 1–23 and 2–24, respectively. Plates were then incubated for 24 hours and processed for the dysferlin quantification and viability cell-based assay. Briefly, each of the 8 plates used in screening were fixed and stained with anti-dysferlin and anti-desmin antibodies. To prevent the discovery of toxic molecules, the number of cells was monitored in parallel by counting the Hoechst stained cells per field and molecule candidates presenting a mortality superior to 50% were excluded.

### Chemical library

Two different small molecule libraries were screened using the 384-well plate format, including 1280 FDA- and EMA-approved drugs from the Prestwick Chemical library and 959 FDA-approved drugs and bioactive compounds from the Selleckchem library. Both libraries were tested at 5 μM.

### Dysferlin quantification and viability cell-based assay

After 24 hours of drug treatment, immunofluorescence was performed on immortalized myoblasts with anti-dysferlin and anti-desmin antibodies and nuclei were visualized with Hoechst staining, as described in immunostaining assay section. Fluorescent labeling was analyzed with a CellInsight CX7 imager (Cellomics Inc). The first channel was used for identification of nuclei, the second for cell segmentation with desmin labeling, and the third one for identification and quantification of dysferlin aggregates. Images were acquired with a 20x objective and were analyzed using a colocalization bioapplication. Using thresholding on the dysferlin staining intensity level, the number of dysferlin-labeled positive cells (cell containing strictly more than one area of dysferlin aggregates) was calculated so as to be less than 10% with the negative control treatment. The total number of cells was determined by counting the Hoechst-stained cells per well, allowing quantification of cell viability by normalization to the negative control.

### Screening data analysis

Screening data analysis was performed using SAW (Sample Assay Warehouse) application (Discngine) connected to Spotfire software (Tibco Software Inc.). The robustness of the assay was evaluated by calculating the Z′ factor on the percentage of dysferlin-labeled positive cells for each plate as follows Z′=1−[3(SDP+SDN)/(MP− MN)]; where MP and MN correspond to the means of the positive (64 nM bafilomycin) and negative (0.1% DMSO) controls, respectively, and SDP and SDN correspond to their standard deviations. Raw data related to cell number per field were normalized to the mean for the negative controls, which is defined as 100%. Hit selection was performed by using the number of standard deviations from the mean in parallel for each readout value (Z-score) calculated per run where all plate data were pooled. Only hits with a Z-score run ⩾5 and that did not decrease the cell number by more than 50%, compared to the 0.1% DMSO condition, were selected for the subsequent validation step. Hits were then re-tested in quadruplicate using the same assay conditions to check that the activity of the molecule was reproducible. Finally, compounds were tested over a range of concentrations (10 nM - 100 µM) for parallel exploration of their efficacy and toxicity, and determination of their half-maximal effective concentration (EC_50_) using nonlinear regression.

### Transient transfection

Healthy immortalized myoblasts were plated on 96-well plates and maintained in growth medium for up to 60% confluence. Myoblasts were transfected by using Lipofectamine RNAiMAX (Invitrogen, United States) according to the manufacturer’s instructions, with siRNA-containing sequences either specific to dysferlin, 5’- TTGGATCAGCTCAGACATATT-3’ or a nonspecific “scrambled” control (Invitrogen, United States). 72 hours after transfection, cells were subjected either to osmotic shock or fixed for dysferlin immunostaining.

### Immunostaining assay

After 24 hours of drug treatment, myoblasts were fixed in 4% paraformaldehyde (10min, room temperature). Immunofluorescence was performed in a phosphate-buffered saline (PBS) solution supplemented with 0.1% Triton X-100 (Thermo Scientific, United States) for permeabilization (5 min, room temperature) and then with 1% bovine serum albumin (BSA; Sigma-Aldrich, United States) for blocking (1 hour, room temperature). Cells were stained for specific markers overnight at 4°C using primary antibodies (listed in Supplementary Table S1). After three successive washes in PBS (5 min, room temperature) labeling was revealed with appropriate fluorophore-conjugated secondary antibodies (listed in Supplementary Table S1) in the dark for 1 h at room temperature, and nuclei were visualized with Hoechst solution (Invitrogen, United States). Following washing steps, coverslips were mounted in fluoromount solution (Thermo Scientific, United States). Images were acquired using a LSM 800 confocal microscope (Zeiss, Germany) with a 63x oil immersion objective and were analyzed with Zen software (Zeiss, Germany).

### Osmotic shock injury assays

Myoblasts were plated in 96-well plates and treated for 24 hours with pharmacological hits. Cells were incubated for 3 hours with a red viability probe (1/3000, Incucyte® Nuclight Rapid Red Dye - Sartorius, Germany) incorporated by the nuclei of living cells, to measure toxicity of molecules (compared with the DMSO negative control) and set osmotic shock-induced mortality at 0%. At the end of the treatment, hypo-osmotic shock was performed by incubating cells with a solution composed of 25% PBS and 75% water, in the presence of a green mortality probe (1/3000, Incucyte® Caspase-3/7 Green Dye - Sartorius, Germany). Green fluorescence was immediately followed every 90 or 120 minutes with an Incucyte® S3 Live-Cell Analysis system (Sartorius, Germany). Images were analyzed with Incucyte software, allowing measurement of the resistance of cells to osmotic shock (green to red ratio) in comparison to the negative control (DMSO).

The two Flexor Digitorum Brevis (FDB) muscles were surgically isolated from euthanized Bla/J and C57BL/6J male mice (6-8 weeks-old, n=6 per conditions), rinsed in D-PBS without Ca^2+^/Mg^2+^ (Gibco, United States) and placed in a digestion solution (collagenase 2,5U/ml in DMEM; Sigma-Aldrich, United States, 11088793001) for 1h30 at 37°C on rotated wheel. The collagenase solution was removed by fiber sedimentation (10 min, room temperature) and the isolated muscle fibers were kept at room temperature in 500µl D-PBS in a 2ml tube. Fibers were treated with bazedoxifene (1µM) or PBS for 40 minutes at room temperature before the induction of osmotic choc by mixing Vol/Vol 100µl of fiber suspensions with 100µl of H_2_O supplemented with 1mM FM1-43 (Life Technologies, United States) in a petri dish. Incorporation of FM1–43 dye into fibers was immediately recorded by a spectral LEICA TCS SP8 confocal microscope (Leica). Tile images were acquired under LASX Software (Leica) using a HC PL Fluotar 10X/0.30 dry objective with a 532 nm laser at 2% power for fluorescence and bright fields. Images of eight acquisition fields were taken every 5 minutes for each mouse, for a total of 30 minutes. A minimum of 84 fibers was examined for each condition. As regards image analysis, a training set of images was done by using the FIJI plugin LabelsToROIs(Waisman *et al*, 2021), to segment and label the fibers. Images were then processed with the Cyto 2 pre-trained segmentation model of Cellpose(Stringer *et al*, 2021; Pachitariu & Stringer, 2022). First, the algorithm was retrained on 20 manually annotated images distinguishing normal and dead fiber cells. Then, each experimental image was segmented by the trained Cyto2, and the position of each fiber was recorded across the 30 minutes of acquisition to monitor the FM1-43 fluorescent intensity. For all conditions, cells already dead at t0 were removed from the analysis. The increase of FM1-43 fluorescent intensity for each fiber of each field for each animal was plotted against the time course of the experiment. Then, a saturation curve fitting was used to measure the coefficient of saturation (a) which is the reflection of the intensity per time unit as a biomarker of the membrane permeability.

### Quantitative PCR

Total RNAs were isolated using the RNeasy Mini extraction kit (Qiagen, Germany) according to the manufacturer’s protocol. A DNase I digestion was performed to avoid genomic DNA amplification. RNA levels and quality were checked using the NanoDrop technology. A total of 500 ng of RNA was used for reverse transcription using the SuperScript III reverse transcription kit (Invitrogen, United States). Quantitative polymerase chain reaction (qPCR) analysis was performed using a QuantStudio 12 K Flex real-time PCR system (Applied Biosystems, United States) and Luminaris color probe qPCR master mixes (Thermo Scientific, United States), following the manufacturers’ instructions. The primer sequence used for *DYSF* gene is commercially available (Applied biosystem, United States): *DYSF* (Hs01002513). Quantification of gene expression was based on the DeltaCt method and normalized to 18S expression (Assay HS_099999).

### Flow cytometry measurement of autophagy flux

Autophagy flux was measured using CYTO-ID® Autophagy Detection Kit (Enzo Life Sciences, United States) according to the manufacturer’s instructions. Briefly, immortalized myoblasts were grown on gelatin 0.1%-coated 6-well plates until 80% confluence and treated for 24 hours with pharmacological compounds. Single-cell suspension was collected after chemical dissociation with trypsin-EDTA (Gibco, United States), centrifuged at 1200 rpm for 5 min, and washed in 1X Assay Buffer. Cells were stained with CYTO-ID® Green Detection Reagent for 30 min at 37°C and protected from light. Cells were washed twice with 1X assay buffer before being assessed by a MACSquant analyzer (Miltenyi Biotec, Germany) with a green fluorescence channel using a 488 nm laser source. A minimum of 30,000 cells were measured per sample. Data were analyzed using FlowJo Software (BD Biosciences, United States). Intact cells were gated in FSC-A vs SSC-A. eGFP median fluorescence intensity (MFI) of all sample was calculated and eGFP-MFI of the DMSO control (= baseline autophagic flux) was substracted to depict the eGFP-MFI shift (=shift of the autophagic flux).

### LC3 puncta formation

Immortalized myoblasts were grown on gelatin-coated 384-well plates and treated with DMSO, bazedoxifene or chloroquine for 24 hours before being fixed with 4% PFA. Immunostaining was performed and LC3 expression was analyzed, employing a 20x objective of the ImageXpress Micro XL System imager (Molecular Devices). A fixed threshold of labeling intensity was chosen for all analyses to detect LC3 puncta, allowing subsequent quantification of their total surface area and integrated intensity for each well (n=9 fields). The total number of cells was determined by counting the Hoechst-stained cells per well (>1000 cells per well), enabling the normalization of LC3 puncta total area or integrated intensity to the nuclei number.

### RFP-GFP-LC3B Tandem Sensor Assay

Immortalized myoblasts were transduced for 24 hours with a Premo™ autophagy tandem sensor RFP-GFP-LC3B (Thermo Scientific, United States) according to the manufacturer’s protocol. Culture medium was replaced and cells were treated for additional 24 hours with 5 μM Bazedoxifene or 0.1% DMSO. After washing with PBS, cells were fixed with 4% formaldehyde and counterstained with Hoechst 33342 solution (Invitrogen, United States). Stained coverslips were mounted with fluoromont solution and images were acquired using a LSM-800 confocal microscope.

### Western blot analysis

Whole-cell lysate of immortalized myoblasts were collected after 24 hours of drug treatment. Proteins were extracted with NP40 lysis buffer (Thermo Scientific, United States) supplemented with 1X Proteases Inhibitors (Complete PIC, Roche, Switzerland). Protein concentration was evaluated using the Pierce BCA Protein Assay Kit (Thermo Scientific, United States) and the absorbance was measured at 562 nm using a CLARIOstar® microplate reader (BMG Labtech, Germany). A total of 20 μg of protein was separated using a 4%–15% Criterion™ XT tris-glycine protein gel (BioRad, United States) and then transferred to nitrocellulose membrane (BioRad, United States) with a Trans-Blot Turbo Transfert system (BioRad, United States) following the manufacturer’s instructions. Membrane was blocked in Odyssey blocking buffer (Li-Cor, United States) for 1h at room temperature and then incubated with primary antibodies (listed in Supplementary Table S1) diluted in blocking buffer from 2h at room temperature to overnight at 4°C. Washing was carried out three times for 10 min at room temperature with TBS + 0.1% Tween 20 (VWR, United States) and the membrane was incubated with appropriate fluorescent secondary antibodies (listed in Supplementary Table S1) in blocking buffer at room temperature for 1h. Washing was carried out, and proteins were detected by fluorescence (Odyssey CLx, Li-Cor, United States) following the manufacturer’s instructions. The expression of proteins was calculated by measuring the intensity of bands using the associated Image Studio^TM^ software. The fold change was calculated after normalization with loading control. All western blotting experiments were performed three or more times, and representative blots are shown.

### Quantification and statistical analysis

Data are presented as means ± SD or SEM for *in vitro* or *ex vivo* experiments, respectively. Statistical analysis were performed using one-way ANOVA followed by Dunnett’s multiple comparisons test, two-way ANOVA with multiple comparisons or unpaired Student’s t-test. Differences were considered significant at P values of \**p*≤0.05, \*\**p*≤ 0.01, \*\*\**p*≤0.001. All graphs were plotted and analyzed using GraphPad Prism Software (v9.2.0).

## ACKNOWLEDGMENTS

The authors thank Marc Peschanski (IStem) for helpful discussions, the research and innovation teams of IStem/CECS for providing expertise and technical support in imaging, bioproduction and screening, the “Imaging and Cytometry Core Facility” of Genethon for technical support, and the GIP Genopole, Evry, and INSERM for the purchase of the equipment. We are grateful to the “Platform for Immortalization of Human Cells” from the Centre of Research in Myology (Institute of Myology, Paris) for providing immortalized myoblasts, and Simone Spuler for providing some of the cellular models used in this study. We would like to thank Marc Bartoli for his expertise in osmotic shock and Tassula Proikas-Cezanne for her advice on autophagy flux. We also thank Celine Leteur and Guillaume Corre for their help in image analysis.

## AUTHOR CONTRIBUTIONS

X.N. and I.R. were responsible for the experimental design and project management. C.B. performed the cell culture experiments, generated the cell banks for screening, developed the screening assay and carried out screening analysis. C.B. and E.P. developed the osmotic shock method and realized functional characterization of the drugs. C.B. studied bazedoxifene’ mechanism of action. J.T. performed the screening and J.P. developed imaging analysis of dysferlin cell based-assay. A.Bi. provided the cellular model. E.P., M.B., N.G. provided technical assistance for cell culture and pharmacological studies. I.R. provided Bla/J mouse model and N.B. performed animal experiments. A.Br. and N.B. developed algorithm quantification of Bla/J fibers resistance to osmotic shock. C.B. prepared the figures. C.B. and X.N. wrote the manuscript. All authors reviewed and edited the paper.

---

**Figure S1.**
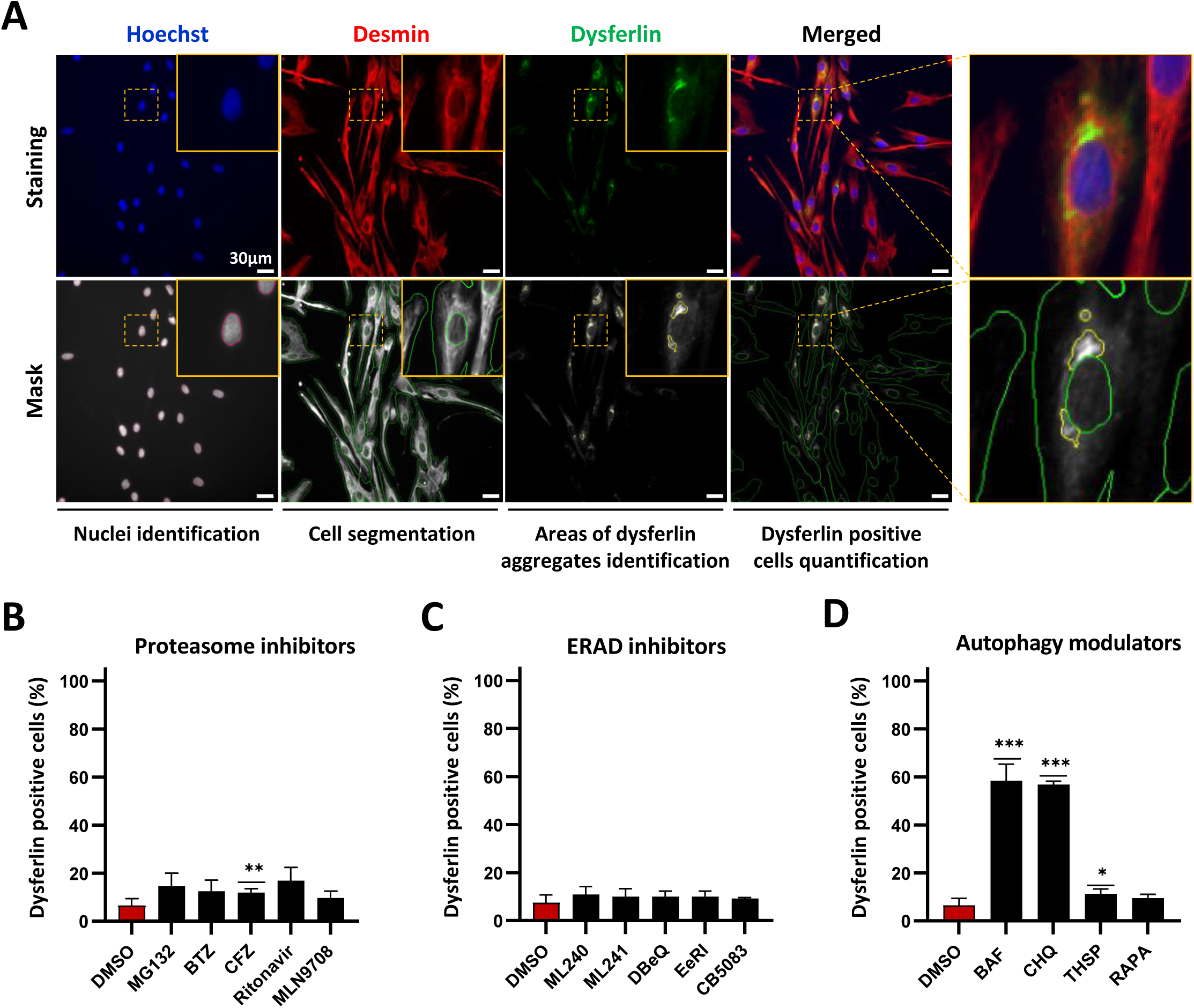
Pilot screen for L1341P dysferlin assay. (**A**) Hoechst, desmin and dysferlin immunostaining in *DYSF*^L1341P^ myoblasts (top panels) and the corresponding detection masks (bottom panels), allowing automated identification of L1341P dysferlin aggregates and quantification of dysferlin-positive cells (>1 aggregate area). Yellow outsets represent higher magnification images. Scale bar = 30µm. (**B-D**) Quantification of dysferlin-positive L1341P myoblasts following treatment with the most effective concentration without toxicity of proteasome inhibitors (**B**), ERAD inhibitors (**C**) or autophagy modulators (**D**) (30µM: CHQ, ML241, ritonavir; 10µM: MG132, BTZ, THSP; 5µM: CFZ, EeRI, ML240, DBeQ, MLN9708; 1µM: CB5083; 300nM: RAPA; 64nM: BAF). Data are shown as the mean ± SD of four replicates. \**p* ≤ 0.05, \*\**p* ≤ 0.01, *** *p* ≤ 0.001 (Brown-Forsythe and Welch ANOVA with Dunnett’s T3 multiple comparisons test). Abbreviations: BTZ, bortezomib; CFZ, carfilzomib; MLN9708, ixazomib; EeRI, eeyarestatin; BAF, bafilomycin; CHQ, choloroquine; THSP, thiostrepton; RAPA, rapamycin.

**Figure S2.**
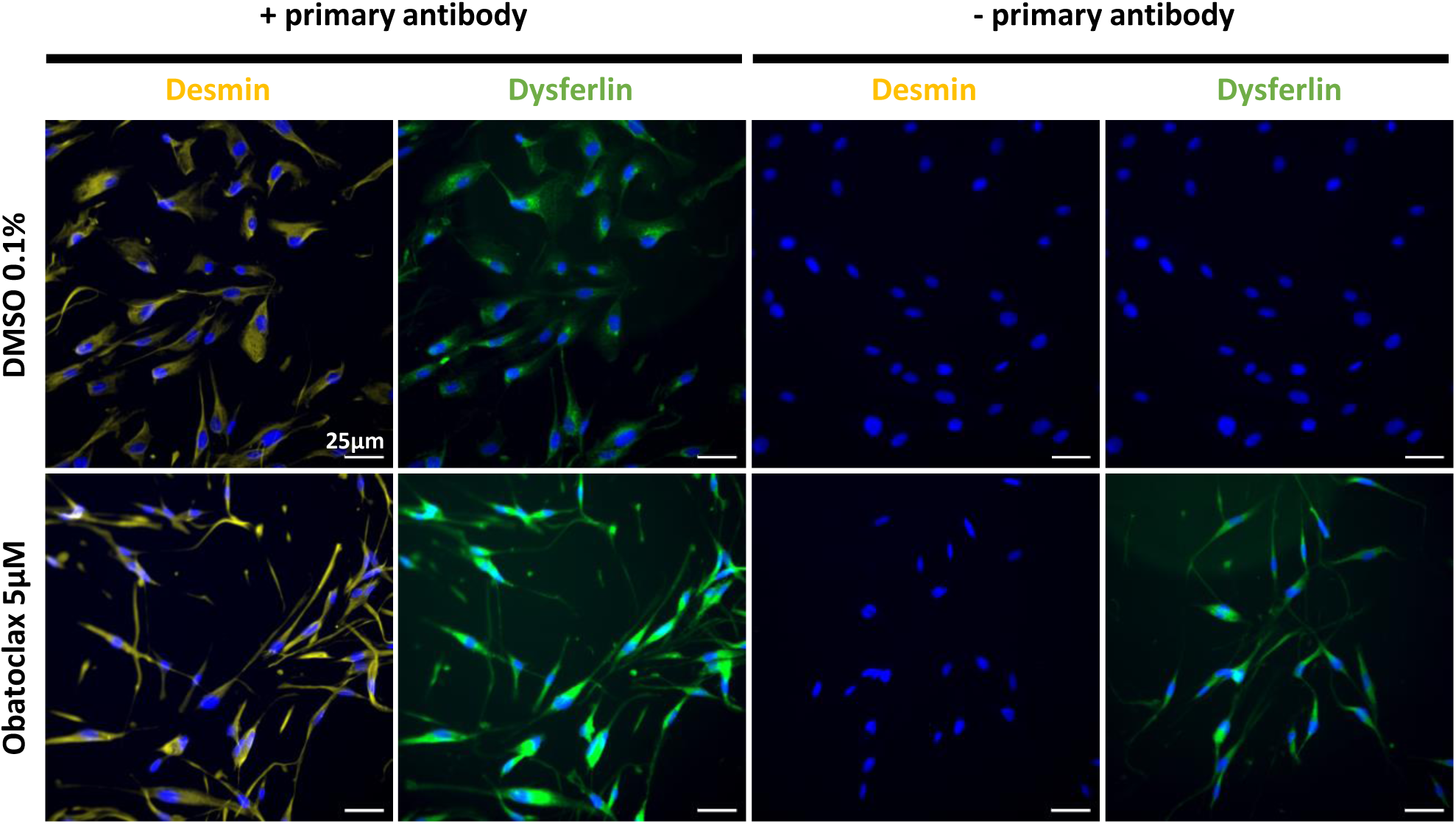
Fluorescence interferences with obatoclax mesylate treatment. Representative images of desmin (yellow) and dysferlin (green) immunostaining in the presence (left panels) or in the absence (right panels) of primary antibodies in *DYSF*^L1341P^ myoblasts treated for 24 hours with 0.1% DMSO or 5 µM obatoclax mesylate. Scale bar = 25 µm.

**Figure S3.**
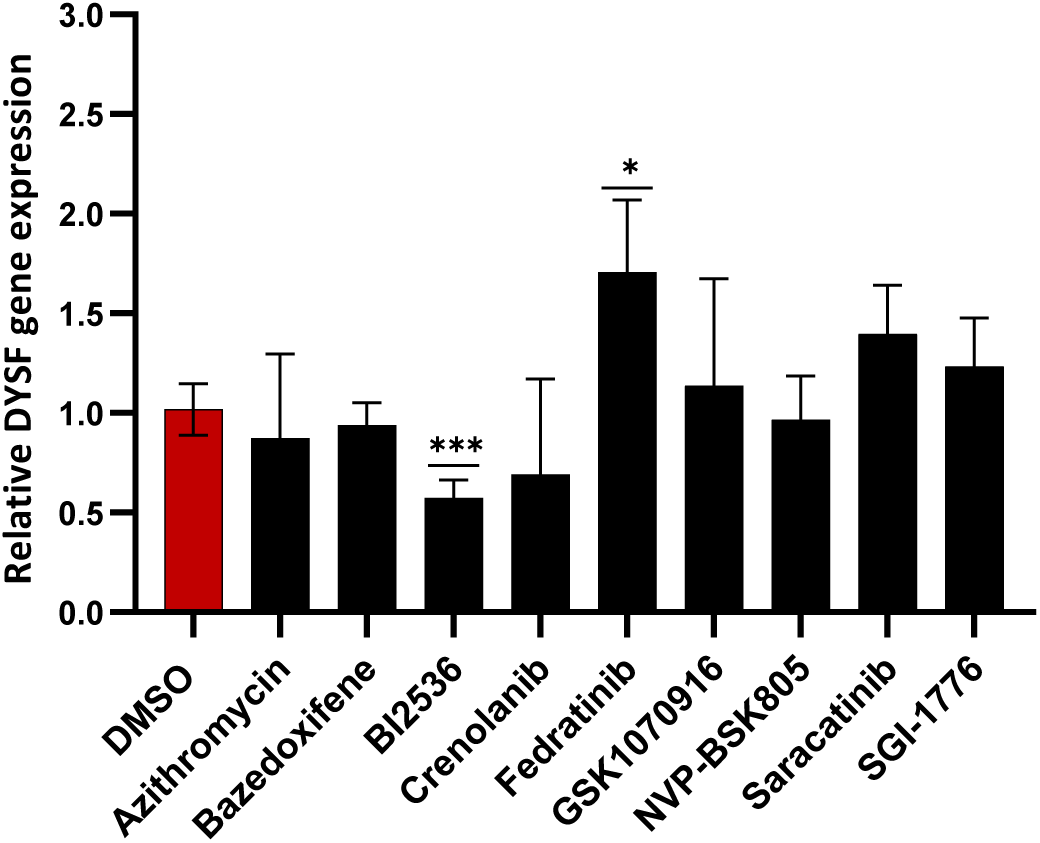
Effect of the nine validated hits on dysferlin gene modulation. Measure of *DYSF* gene expression by qPCR in *DYSF*^L1341P^ myoblasts treated for 24 hours with 0.1% DMSO or the 9 validated hits at their maximum effective dose (10µM: azithromycin; 5µM: bazedoxifene, crenolanib, GSK1070916, saracatinib; 2µM: BI2536, fedratinib, *NVP*-BSK805, SGI-1776). Gene expression is normalized to DMSO-treated cells. Data are shown as the mean of three independent experiments ± SD. \**p* ≤ 0.05, \*\*\**p* ≤ 0.001 (Brown-Forsythe and Welch ANOVA with Dunnett’s T3 multiple comparisons test).

**Figure S4.**
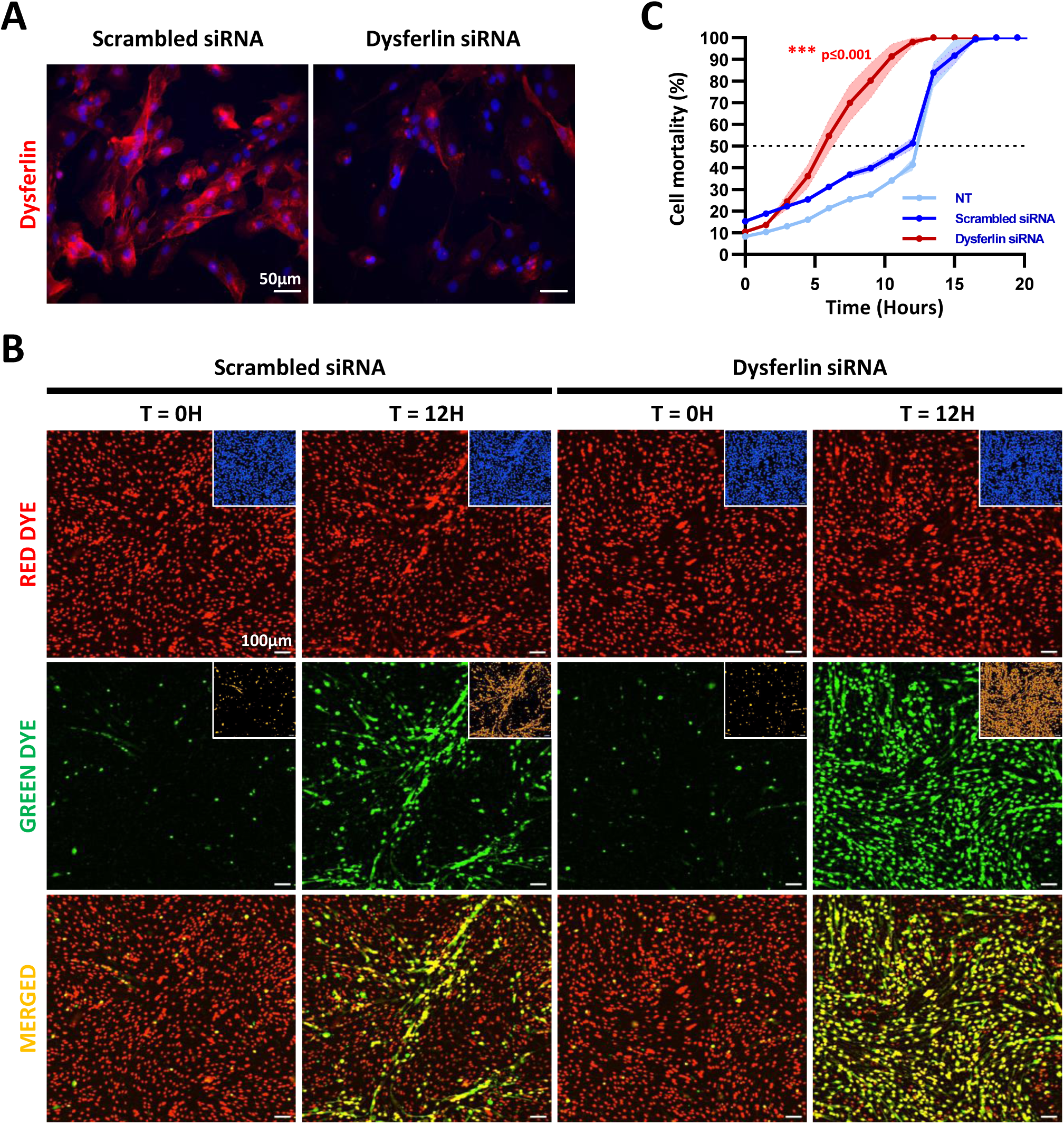
Dysferlin deficient myoblasts are more susceptible to membrane damage following hypo-osmotic shock. (**A**) Representative images of dysferlin immunostaining (red) of healthy immortalized myoblasts treated for 72 hours with scrambled or dysferlin siRNA. Scale bar = 50µm. (**B**) Live labeling with red dye, for nuclear staining, and green dye, for caspase3/7 apoptosis, of healthy immortalized myoblasts subjected to hypo-osmotic shock after 72 hours of transfection with scrambled (left panels) or dysferlin (right panels) siRNA. Representative images of labeling and corresponding detection mask pictures, on the top right of each image, for automated quantification of cell mortality (green/red ratio) at the beginning of osmotic shock (T=0H) or when 50% mortality is achieved with scrambled siRNA treatment (T=12H). Scale bar = 100µm. (**C**) Quantification of cell mortality after induction of osmotic shock in healthy myoblasts treated for 72 hours with scrambled (blue) or dysferlin (red) siRNA. Each point represents the mean ± SD of three replicates on a representative control cell line. \*\*\**p* ≤ 0.001 (One-way ANOVA with Dunnett’s multiple comparisons test).

**Figure S5.**
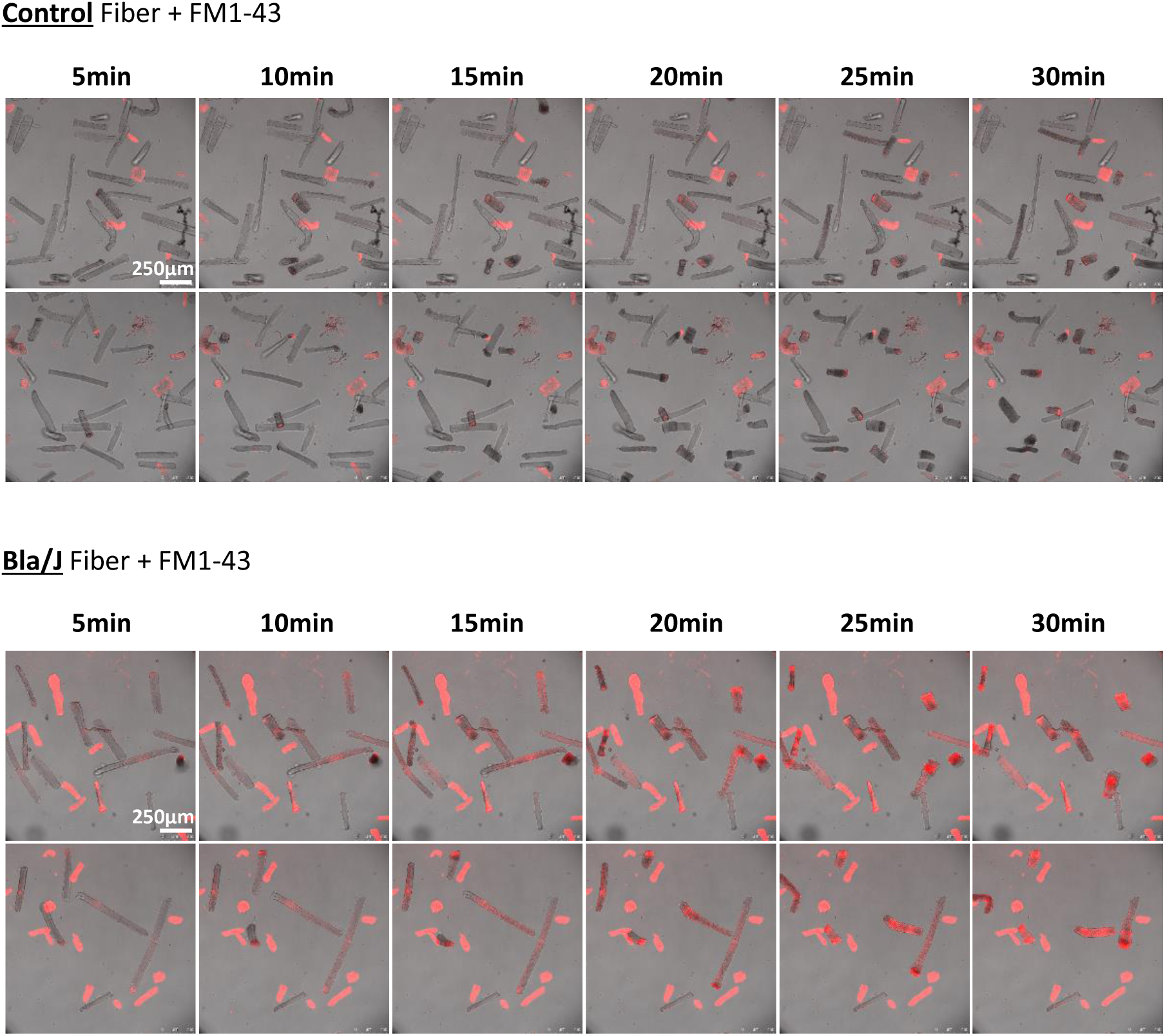
Defective resealing of membranes in dysferlin-null myofibers from Bla/J mice. Representative images of FM1-43 dye accumulation in myofibers isolated from control (upper panel) or Bla/J (lower panel) animal and subjected to hypo-osmotic shock. Fluorescence was monitored every 5 minutes for a total duration of 30 minutes. Scale bar = 250 µm.

**Figure S6.**
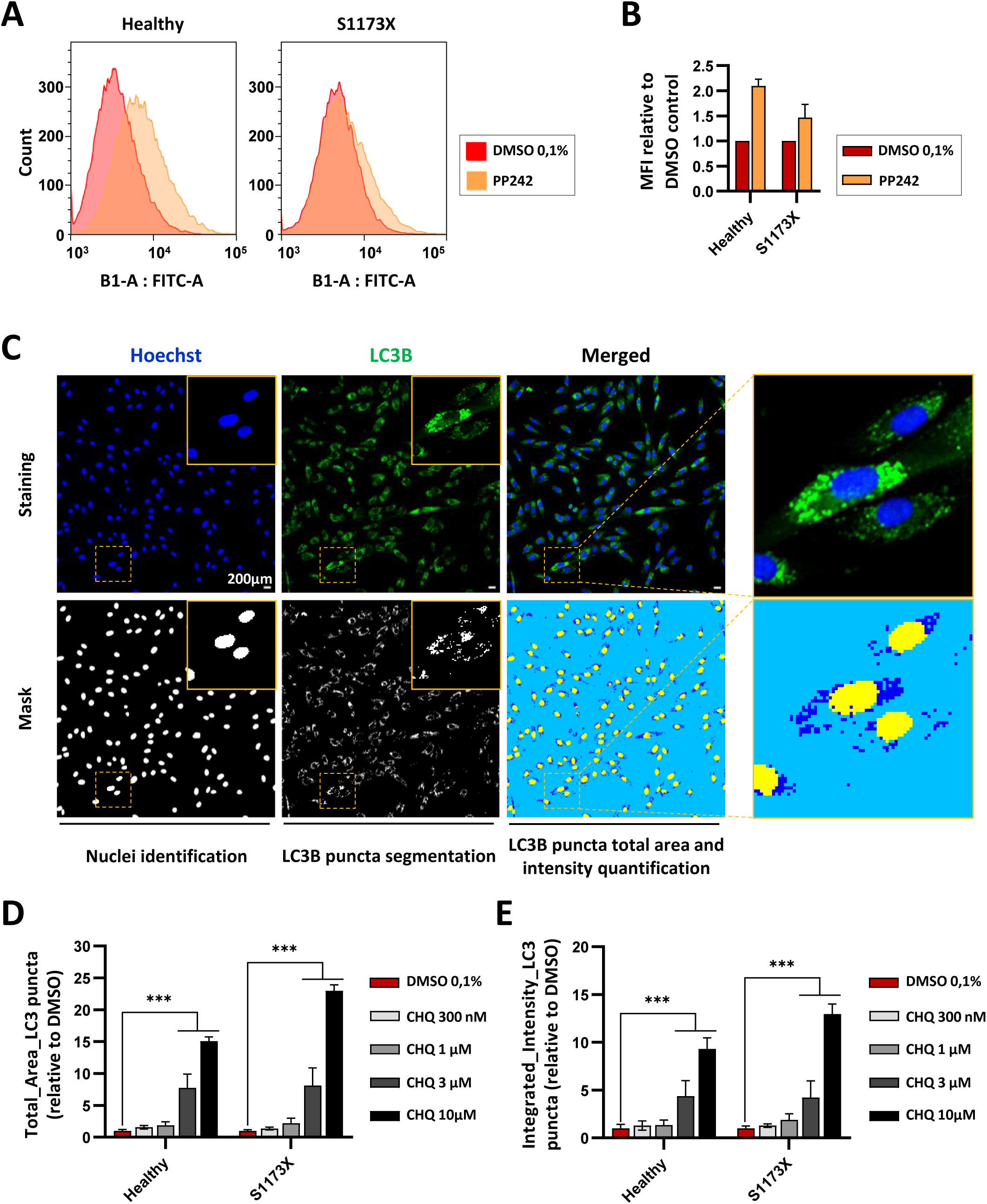
Variations of autophagy flux after treatment with autophagy modulators. (**A**) Representative FACS plots showing eGFP fluorescence detected using a CYTO-ID® Autophagy detection kit in healthy and *DYSF*^S1173X^ myoblasts, treated with or without PP242 (10µM) for 24 hours. (**B**) Median fluorescence intensity (MFI) was assessed for each condition and normalized to DMSO control. Data are shown as the mean ± SD (n=2). (**C**) Representative images showing Hoechst (blue) and LC3B (green) immunostaining in *DYSF*^S1173X^ immortalized myoblasts treated with 10µM chloroquine for 24 hours (top panels) and the corresponding detection masks (bottom panels), resulting from the implementation of an automated detection and quantification of the LC3 puncta algorithm in the MetaXpress software. Yellow outsets represent higher magnification images. Scale bar = 200 µm. (**D-E**) Quantification of the relative LC3 puncta average total area (**D**) and integrated intensity (**E**) in healthy and *DYSF*^S1173X^ myoblasts following 24 hours treatment with increasing concentrations of chloroquine. Each bar represents the mean ± SD (n=4 or more) of a representative experiment over three independent experiments. \*\*\**p*≤0.001 (2way ANOVA with multiple comparisons). Abbreviation: CHQ, chloroquine.

**Supplementary Table S1.** List of antibodies (related to Materials and Methods).

| Application | Antibodies | Supplier | Reference | Dilution |
| --- | --- | --- | --- | --- |
| Immunostaining | DYSFERLIN | Abcam | Ab124684 | 1 :200 |
|  | DESMIN | R&D | AF3844 | 1 :200 |
|  | KDEL | EnzoLife | ADI-SPA-827-D/F | 1 :200 |
|  | LC3B | Cell Signaling | 2775S | 1 :1 000 |
|  | Donkey anti-rabbit<br>AF488 | Invitrogen | A-21206 | 1 :1 000 |
|  | Donkey anti-mouse<br>AF555 | Invitrogen | A-31570 | 1 :1 000 |
|  | Donkey anti-goat<br>AF555 | Invitrogen | A-21432 | 1 :1 000 |
|  | Donkey anti-goat<br>AF647 | Invitrogen | A-21447 | 1 :1 000 |
| Western Blot | LC3B | Novus Biologicals | NB600-1384 | 1 :500 |
|  | β-ACTIN | Li-Cor | 926-42210 | 1 :1 000 |
|  | IRDye® 800CW<br>Donkey anti-rabbit | Li-Cor | 926-32213 | 1 :5 000 |
|  | IRDye® 680RD<br>Donkey anti-Mouse | Li-Cor | 926-68072 | 1 :10 000 |

